# Magnetic resonance control of reaction yields through genetically-encoded protein:flavin spin-correlated radicals in a live animal

**DOI:** 10.1101/2025.02.27.640669

**Authors:** Shaun C. Burd, Nahal Bagheri, Maria Ingaramo, Alec F. Condon, Samsuzzoha Mondal, Dara P. Dowlatshahi, Jacob A. Summers, Srijit Mukherjee, Andrew G. York, Soichi Wakatsuki, Steven G. Boxer, Mark Kasevich

## Abstract

Radio-frequency (RF) magnetic fields can influence reactions involving spin-correlated radical pairs. This provides a mechanism by which RF fields can influence living systems at the biomolecular level. Here we report the modification of the emission of various red fluorescent proteins (RFPs), in the presence of a flavin cofactor, induced by a combination of static and RF magnetic fields. Resonance features in the protein fluorescence intensity were observed near the electron spin resonance frequency at the corresponding static magnetic field strength. This effect was measured at room temperature both in vitro and in the nematode *C. elegans*, genetically modified to express the RFP mScarlet. These observations suggest that the magnetic field effects measured in RFP-flavin systems are due to quantum-correlated radical pairs. Our experiments demonstrate that RF magnetic fields can influence dynamics of reactions involving RFPs in biologically relevant conditions, and even within a living animal. These results have implications for the development of a new class of genetic tools based on RF manipulation of genetically-encoded quantum systems.

---

Remote regulation of biomolecular reactions in living organisms could enable new scientific advances and potential therapies. Radio-frequency (RF) magnetic fields are ideal for this purpose as biological tissues are essentially transparent to this part of the electromagnetic spectrum. Here we demonstrate control of reaction yields using RF magnetic fields in a protein system genetically encoded in a living animal.

Although energy scales associated with typical static and RF magnetic fields are orders of magnitude below thermal fluctuations in biologically relevant conditions, RF fields can alter reaction yields in chemical and biochemical systems involving spin-correlated radical pairs (SCRPs) - a phenomenon known as reaction-yield detected magnetic resonance (RYDMR) (*1–3*). This is observed when a time-varying magnetic field is applied near the electron spin resonance (ESR) frequency *f*_ESR_ = *gμ*_0_*B*_0_/ℎ in a static magnetic field *B*_0_, where *g* is the electron g factor, *μ*_0_ is the

Bohr magneton, and ℎ is Planck’s constant. Moreover, observations that RF fields can influence biological processes in animals, including the magnetic sense of birds (*4*), have also been attributed to radical-pair dynamics (*5*). However, the detailed mechanism behind these effects at the protein level remains uncertain (*6*). Previous studies of radical pairs in protein systems (*7–14*) have focused on the effects of static magnetic field effects (MFEs), with the notable exception of chemically-modified photosynthetic reaction centers, where the influence of time-varying magnetic fields has been investigated extensively in vitro (*15–17*).

Here, we show that RF fields applied near the ESR frequency can modulate reaction yields in a system consisting of a red fluorescent protein (RFP) paired with a photoproduct of flavin mononucleotide (FMN) (Fig. 1). Reaction yields in this system are inferred from changes in the RFP fluorescence intensity induced by the RF field. RFPs have become essential tools in biological and photophysical research (*18, 19*) and FMN is a naturally occurring molecule in cells. We present in vitro results showing RYDMR for various RFPs including mScarlet, mScarlet-I (*20*), mCherry (*21*), and mCherry-XL (*22*) together with FMN, and in vivo measurements in an animal, a transgenic *C. elegans* nematode, engineered to express mScarlet in all cells (*23, 24*).

**Figure 1:**
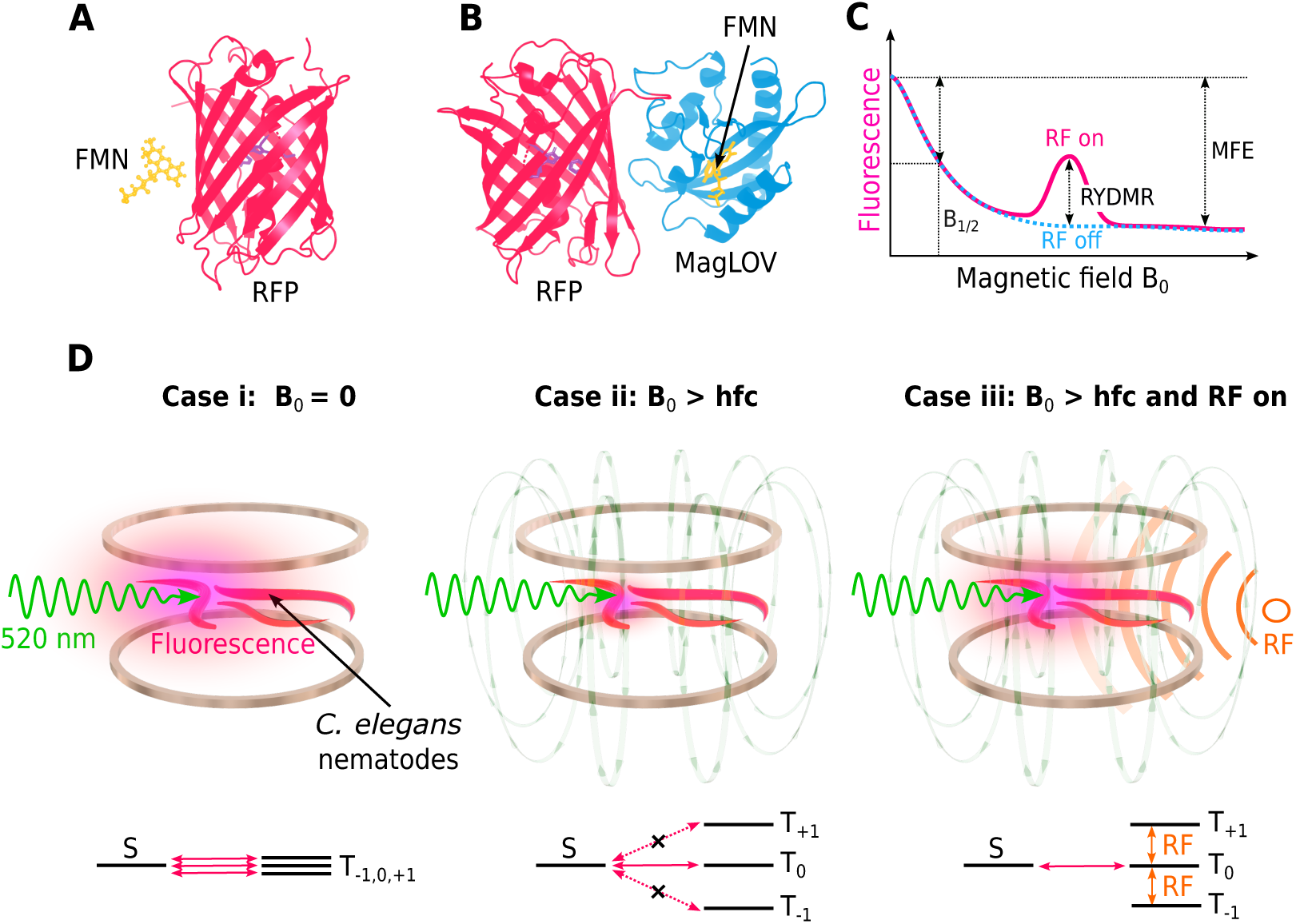
Modulation of fluorescence from red fluorescent proteins (RFPs) in a fluorescent protein: flavin system using radio-frequency magnetic fields. (**A**) Structure of mScarlet (PDB ID: 5LK4) together with an illustration of flavin mononucleotide (FMN). (**B**) Rendering of mScarlet-MagLOV fusion protein (PDB IDs: 5LK4 and 7PGY). (**C**) Schematic of predicted RFP fluorescence intensity as a function of the static magnetic field strength *B*_0_, with continuous RF applied at the ESR frequency. *B*_1/2_ is the magnetic field at which the fluorescence has decreased by half of the maximum magnetic field effect (MFE). The increase in the fluorescence resulting from the RF is the reaction yield detected magnetic resonance (RYDMR). (**D**) Illustrations of *C. elegans* nematodes expressing RFP fluorescing under green light excitation are shown in the upper row. Pre-excitation with blue light (not shown) is used for flavin photoproduct generation from endogenous flavin prior to green light excitation. Helmholtz coils generate a magnetic field *B*_0_. Corresponding singlet (*S*) and triplet (*T*) energy levels from a putative spin-correlated radical-pair are shown beneath. With *B*_0_ = 0, *S* − *T* mixing can occur between *S* and all triplet sublevels (Case i). When *B*_0_ exceeds the nuclear hyperfine constants (hfc) that drive *S* − *T* mixing (Case ii), the *T*_±1_ sublevels become energetically isolated and only *T*_0_ can be converted to *S* efficiently, resulting in a reduction of RFP fluorescence. Application of an RF magnetic field at the ESR frequency redistributes population between the triplet sublevels, resulting in increased fluorescence (Case iii).

A custom apparatus (Fig. S1) enables widefield fluorescence imaging of biological samples expressing FPs or purified proteins in aqueous solution inside an RF resonator. We typically use a bridged loop-gap resonator (BLGR) (*25, 26*) with a resonance frequency *f*_res_ ∼ 450 MHz and a quality factor of ∼ 600 (see SI). Static fields from 0 to ∼ 30 mT parallel to (*B*_0 ∥_) or perpendicular to (*B*_0 ⊥_) the RF magnetic field *B*_1_ can be generated using two sets of Helmholtz coils. For photoexcitation, lasers at 440 nm (blue) and 520 nm (green) are applied to the sample. We typically apply an initial ∼10 s duration 440 nm photosensitization pulse that creates an as-yet not characterized photoproduct of the flavin, and subsequently apply 520 nm excitation for the remainder of the experiment (see SI). Fluorescence from the sample passes through a 650 nm long-pass filter and is imaged using a camera. The filter cuts off residual FMN fluorescence and serves to block scattered excitation light. Aqueous samples are placed inside a 3 mm inner diameter quartz tube, while *C. elegans* samples are placed on an agarose substrate on a fused-silica support, both near the center of the RF resonator.

Measurements of the fluorescence from an aqueous solution of purified mScarlet-I together with FMN pre-excited with blue light are shown in Fig. 2 A. Increasing the magnetic field *B*_0_ initially results in a decrease in fluorescence, as observed in a recent report (*27*). Near *B*_0_ = 15.9 mT — the field required for ESR at the RF frequency *f*_RF_ = 447 MHz — there is a resonance feature in the fluorescence. Increasing the RF magnetic field amplitude *B*_1_ results in a linear increase in the peak of the resonance and also an increase in the resonance full width at half maximum (FWHM) (Fig. 2 B and C) as estimated by using a least-squares fit of the sum of two Lorentzian functions to the data in Fig. 2 A (see SI). These features are consistent with RYDMR theory for the regime where *B*_1_ is weak compared to the interactions driving spin-state mixing (*3, 28*). The fit is also used to estimate the MFE, quantified by the fractional reduction in the fluorescence intensity of the FP when the static magnetic field is applied, and *B*_1/2_ — the field at which the fluorescence has decreased by half the estimated maximum possible value. The independence of both the estimated maximum MFE and *B*_1/2_ values from the RF excitation strength *B*_1_ suggests that the system is only weakly perturbed by the RF, as expected in the limit *B*_1_ ≪ *B*_0_ (Fig. 2 D and E). To measure the dynamic response of the resonance fluorescence to changes in *B*_1_, the RF field was stepped between a given value of *B*_1_ and zero within 500 *μ*s as shown in Fig. 2 F, with optical excitation as indicated in the figure. The dynamic response was inferred by fitting a first-order exponential decay model to the data (Fig. 2 G, see SI), to estimate the time constant (*τ*) of the response and the RYDMR amplitude (Figs. 2 H and I). We find *τ* to be largely independent of *B*_1_ (Fig. 2 H), for a fixed value of the 520 nm excitation intensity (*I*_520_). Increasing *I*_520_ from 0.5 W/cm^2^ to 3.8 W/cm^2^ results in a 2-fold increase in the magnetic resonance amplitude and a concomitant reduction in *τ* from ∼ 2.5 s to ∼ 1.5 s (Figs. 2H and I). As *I*_520_ increases, the increase in the RYDMR amplitude becomes smaller, suggesting that the optical interaction becomes saturated. Increasing the 440 nm preexcitation intensity *I*_440_, with fixed *I*_520_, results in a negligible increase in the RYDMR amplitude beyond *I*_440_ ∼ 1 W/cm^2^.

**Figure 2:**
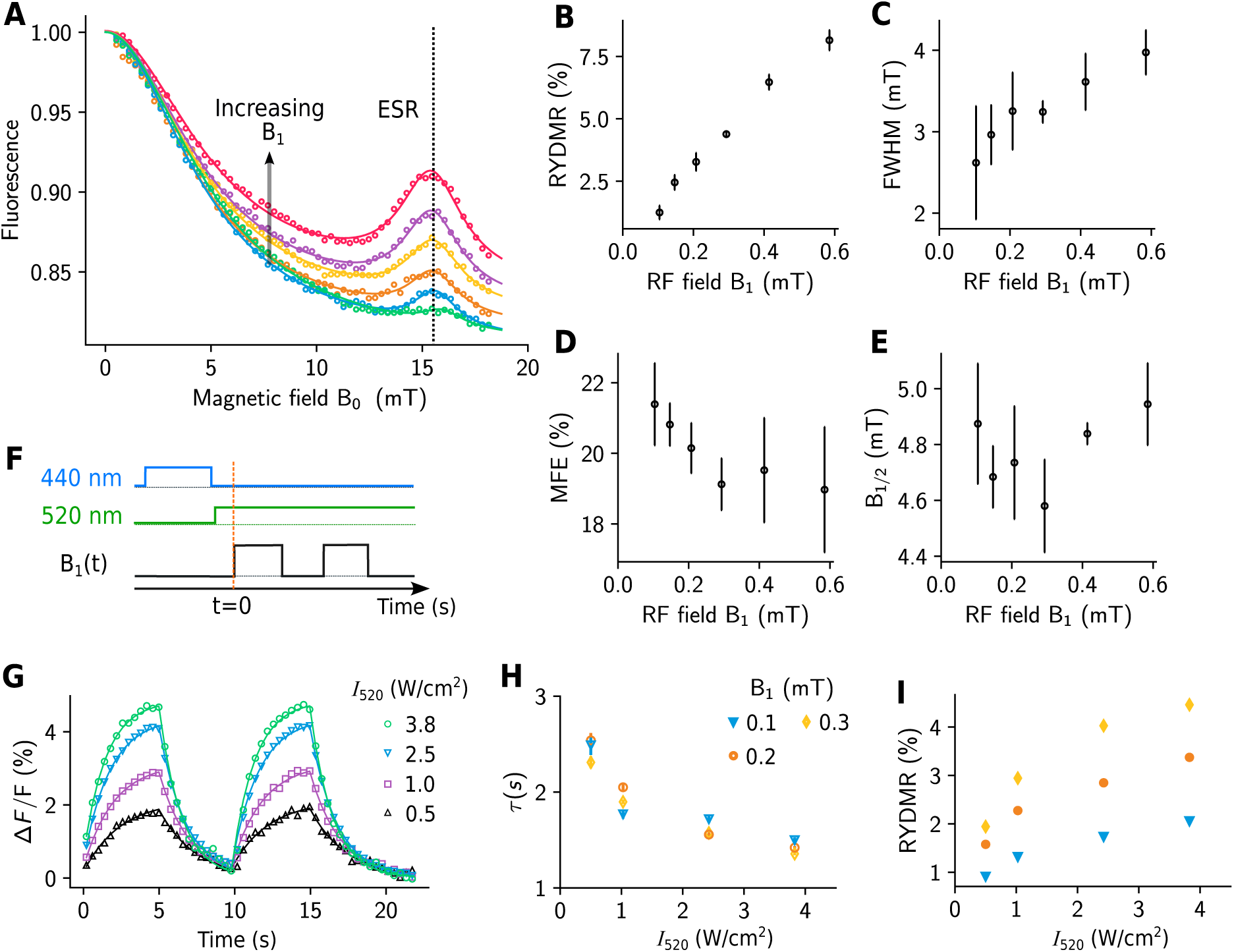
RYDMR from an aqueous solution of purified mScarlet-I and FMN. (**A**) Fractional changes in mScarlet-I fluorescence at various values of the DC field *B*_0 ⊥_ in the presence of a 447 MHz RF field with increasing values of *B*_1_ ranging from 0.07 to 0.37 mT. Solid lines are fits to the data as described in the main text and SI. (**B**) to (**E**), Resonance amplitude (RYDMR), full width at half maximum (FWHM), magnetic field effect (MFE), and *B*_1/2_ derived from the fit parameters (see Fig. 1 C and SI). Error bars indicate the estimated standard errors from three technically repeated experiments. The vertical dashed line in **A** indicates the ESR magnetic field (15.9 mT) corresponding to 447 MHz. (**F**) Timing diagram (not to scale) for measuring changes in fluorescence resulting from step changes in the RF field (between a specified value of B_1_ and 0 mT) with *B*_0 ⊥_ = 15.9 mT at various 520 nm intensities (*I*_520_). (**G**) Time traces showing fractional changes in fluorescence (Δ*F*/*F*) with B_1_ = 0.3 mT from t = 0 - 5 s and 10 - 15 s and B_1_ = 0 elsewhere. Solid lines are fits to experimental data (see SI) to estimate resonance fluorescence time constants (**H**) and amplitudes (**I**) at various values of *I*_520_ and *B*_1_. For all experiments, concentrations of mScarlet-I and FMN are 50 *μ*M and 350 *μ*M respectively. Experiments were conducted at 21.5^◦^ C.

The value of *B*_0_ at which the resonance feature occurs was investigated by testing samples in RF resonators with different center frequencies. The measured values of the line-center field *B*_0res_ at different RF frequencies are plotted in Fig. 3 A, showing good agreement with the theoretical prediction for the ESR. Furthermore, if the static field is parallel to the RF field, we do not observe any obvious resonance features (Fig. 3 B) as expected from the ESR transition selection rules.

**Figure 3:**
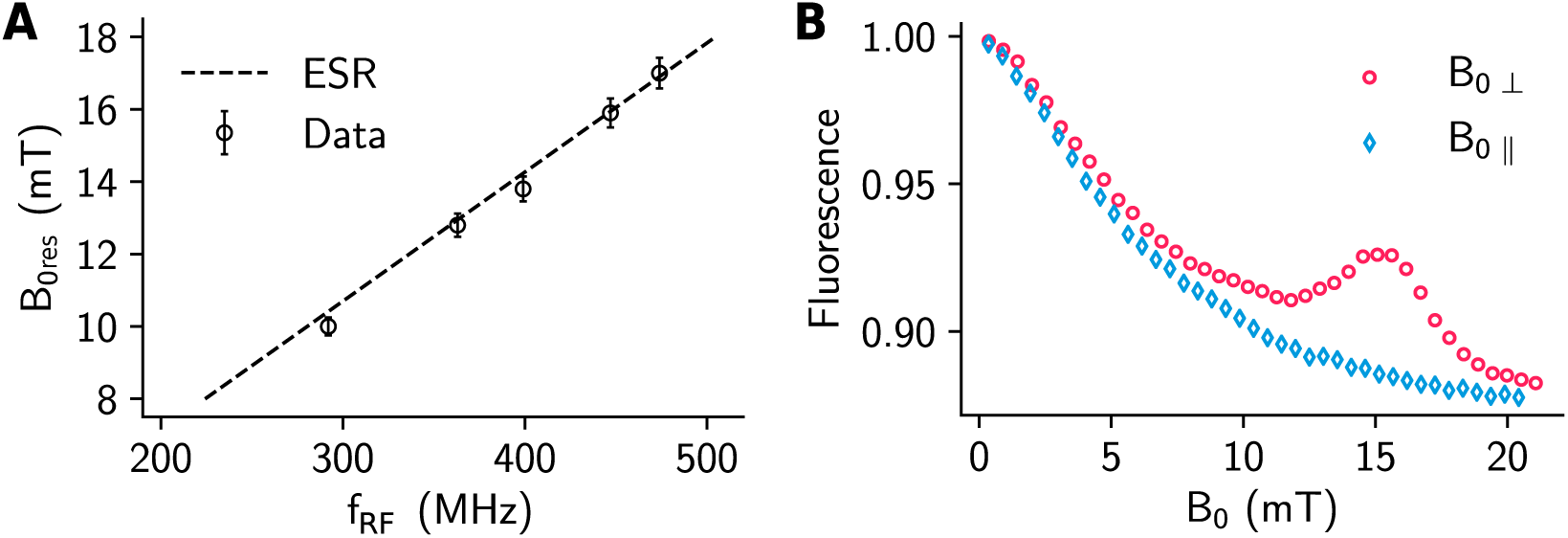
Electron spin resonance. (**A**) Measured resonance magnetic field *B*_0res_ at various values of the RF drive frequency. The dashed line is the ESR magnetic field strength as a function of the RF frequency calculated from *B*_0_ = ℎ *f*_RF_/*μ*_0_*g*. Error bars indicate the uncertainty in the magnetic field calibration. (**B**) Measured fluorescence at various values of the static field, parallel (B_0 ∥_) or perpendicular (B_0 ⊥_) to the RF field B_1_.

We have observed MFEs and RYDMR from multiple RFPs together with FMN. The largest MFEs are measured for mScarlet and its variants (mScarlet-I, mScarlet3) at the ∼ 20 % level (Fig. 2 and SI) and the smallest for mCherry at ∼ 1.5 %. Similarly, the largest RYDMR features are measured in mScarlet and its variants reaching almost 10% increase in fluorescence (Fig. 2 A, B). Measurements for mCherry, and two of it variants show that mutations can have an effect on MFE and RYDMR characteristics (see SI).

Our setup enables detection of magnetic resonance dependent fluorescence in the widefield, as the BLGR geometry inherently provides a near-uniform oscillating magnetic field amplitude *B*_1_ over the area enclosed by the resonator (*25*), and the Helmholtz coils provide *B*_0_ uniformity better than 0.3 % over a volume of ∼ (1.5 cm)^3^ (see SI). These features enable measurement of RYDMR from multiple *C. elegans* nematodes simultaneously (Fig. 4). Spatially-resolved RYDMR measurements are obtained by measuring the fluorescence from a specified region of interest of the image, while the magnetic field *B*_0_ is varied. Mapping out the RYDMR amplitude over the entire image (Fig. 4 C and F) shows that measurement of RYDMR is spatially correlated with mScarlet fluorescence originating from the nematodes. In comparison to the 20 % level MFEs measured in vitro (Fig. 2 **A** and **D**), the largest MFEs measured in *C. elegans* are around 4 %. One possible explanation might be the lower levels of endogenous free flavin cofactor in the animals. However, the magnetic field at which the RYDMR resonance occurs, *B*_0res_, is largely insensitive to the cofactor concentration.

**Figure 4:**
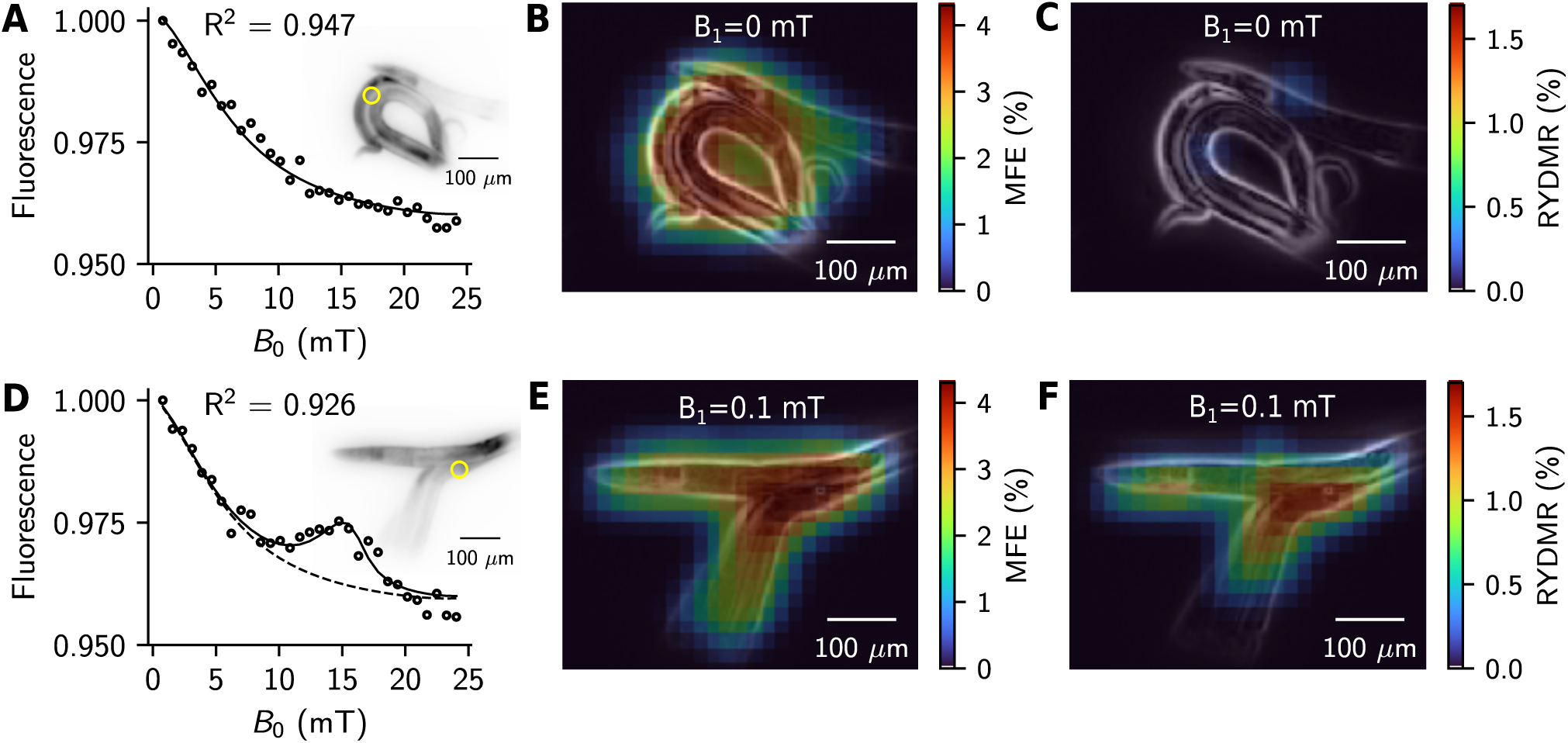
Magnetic field effects and RYDMR in *C. elegans* expressing mScarlet in all cells. (**A**) to (**C**) Experimental data where the RF is off, *B*_1_ = 0 mT. (**D**) to (**F**) *B*_1_ = 0.1 mT at 445 MHz. (**A**) and (**D**) Fluorescence from *C. elegans* at various values of the static magnetic field *B*_0_. The insets show grayscale mScarlet fluorescence images of the nematodes used for the experiments. Fluorescence from the regions within the yellow circles in the insets is plotted in the respective main panels. The solid line is a fit to the data (black circles) as described in the SI. The dashed line indicates the estimated fluorescence with no RF. (**B**) and (**E**) MFE distribution overlaid on an edge map (EM) derived from the fluorescence images of the nematodes shown in **A** and **D**. (**C**) and (**F**) RYDMR distribution overlaid on the EM of the nematodes.

We have also investigated the engineered flavin-binding fluorescent protein MagLOV (*27*) and an mScarlet-MagLOV fusion construct which possibly show reduced cofactor concentration dependence of the MFE and RYDMR amplitude as the flavin is collocated with the RFP (Fig. 1 B). Measurements of RYDMR from *E. coli* colonies expressing these engineered proteins are given in the SI.

The characteristics of the magnetic resonance features in the RFP-flavin system suggest that a spin-correlated radical pair is responsible for the observed magnetic effects. A proposed simplified reaction scheme is illustrated in Fig. S3. Following initial photoexcitation of the FMN cofactor near its absorption peak (∼ 440 nm), subsequent photoexcitation with green light results in the formation of a SCRP. This is likely due to electron transfer or hydrogen atom abstraction with an amino acid(s) on the surface of the RFP by the excited flavin or its photoproduct, as shown in early photochemically-induced dynamic nuclear polarization (photoCIDNP) studies of amino acids:flavin systems (*29, 30*). Note that the physics underlying CIDNP is the same RP mechanism that underlies the magnetic field effects. At low external magnetic fields, hyperfine coupling to magnetic nuclei in the radical partners drives coherent singlet (*S*) to triplet (*T*) interconversion of the two-electron spin state of the radical pair (Fig. 1 D). Singlet RPs are returned to their molecular ground states through a reverse reaction. Triplet RPs can return to the ground state either via conversion to *S* through coherent spin-state mixing or through incoherent spin relaxation on much slower time scales. In this model, we assume that the ground state of the RFP is the only form of the RFP that can be excited to a state that fluoresces appreciably upon illumination with green light. The resulting fluorescence is sensitive to the spin state of the radical pair, as it affects the population of the ground state of the RFP. This could occur by direct involvement of the RFP chromophore or indirectly by interaction of the fluorescing state of RFP with one partner of the RP, e.g. a free radical on the surface of the RFP. In the absence of an applied magnetic field, the RP triplet sublevels *T*_−1_, *T*_0_, and *T*_1_ are close to degeneracy and coherent interconversion can occur between *S* and all triplet sublevels (Fig. 1 D Case i). If an external magnetic field with flux density *B*_0_ is applied, the energy splitting between the triplet sublevels increases due to the Zeeman effect. If *B*_0_ is increased such that Zeeman splitting exceeds the hyperfine coupling driving the coherent spin-state mixing, the *T*_±1_ levels become energetically isolated and coherent interconversion can only occur appreciably between the *S* and *T*_0_ states (Fig. 1 D Case ii). Thus, for a radical pair born in the triplet state, an external magnetic field will reduce the steady state population of ground-state RFPs resulting in a reduction of measurable fluorescence (Fig 1 D Case ii). Note that there is a smooth decrease in fluorescence as *B*_0_ increases, so there is no evidence for residual exchange coupling in the RP. If an RF magnetic field *B*(*t*) = *B*_1_ sin(*f*_RF_*t*), is applied in a direction orthogonal to *B*_0_, where the frequency *f*_RF_ matches the electron Larmor frequency, transitions between *T*_0_ and *T*_±1_ can occur. The coupling between triplet states by the RF field enables population in the *T*_±1_ states to access *T*_0_ and consequently *S* (Fig 1 D, case iii), resulting in an increase in fluorescence when the RF field frequency is near the electron Larmor frequency (Fig 1 D). The resonance fluorescence linewidth can be used to give an estimate of the coherence lifetime of the radical pair. The narrowest measured FWHM of 2.7(7) mT (Fig. 2 C) corresponds to a frequency linewidth of Δ*v* = 72 MHz and a coherence time of 1/*π*Δ*v* = 4(1) ns — potentially enabling radical-pair-based biosensing applications (*31*).

Our work also suggests that RFPs could serve as genetically-encoded fluorophores for various magnetic imaging modalities that have traditionally relied on chemical agents unsuitable for biological applications (*32*). FP mutants with enhanced magnetic resonance characteristics and field sensitivity could be generated using directed evolution approaches (*27*).

We have demonstrated that RF magnetic fields can modulate reaction dynamics in a protein system at room temperature in vitro and in a living animal. This opens up opportunities for designing RF-controllable genetically-encoded systems for regulation of other biomolecular processes such as cell signaling or gene expression. We anticipate that engineering genetically-encoded quantum entangled states in biological systems could enable interfacing living organisms with emerging quantum technologies.

## Acknowledgments

We thank the laboratory of Prof. Kang Shen for assistance preparing *C. elegans* samples. We thank Prof. Kang Shen, Dr. Bing Xu, Dr. Callista S. Yee, Dr. Eric I. Rosenthal, and Joshua L. Reynolds for helpful discussions. We thank Prof. Ralph Jimenez and Dr. Nancy Douglas for providing plasmids of RFP mutants.

## Funding

We acknowledge funding from the US Department of Energy, BER FWPs 100878 and 100882 (M.K. and S.W.); NIH grant R35GM118044 (S.G.B.); N.B. acknowledges support from the Stanford Bio-X Bowes Graduate Student Fellowship.

## Author contributions

Conceptualization: S.C.B., N.B., M.K., S.G.B., S.W., M.I., A.G.Y., Methodology: S.C.B., N.B., M.K., S.G.B., S.W., Investigation: S.C.B., N.B., M.I., A.F.C., S.M., D.P.D., J.A.S., Visualization: S.C.B., N.B., M.K., Funding acquisition: M.K., S.G.B., S.W., Project administration: M.K., S.G.B., S.W., Supervision: S.C.B., M.K., S.G.B., S.W., Writing – original draft: S.C.B., N.B., Writing – review editing: All authors

## Competing interests

A.G.Y. and M.I. are listed as inventors in U.S. Provisional Patent Application No. 63/568,263, entitled “MUTANT ASLOV2 DOMAINS AND USES THEREOF.”

## Supplementary Materials

## 1 Materials and Methods

### 1.1 Experimental setup

The experimental setup is illustrated in Fig. S1. Static magnetic fields along the *x* (*B*_0 ∥_) and *z* (*B*_0 ⊥_) directions are generated using pairs of coils in the Helmholtz configuration. Programmable power supplies (Keysight E36233A and E36155A) enable generation of fields ranging from 0 to 25 mT along *x* and 0 to 30 mT along *z*. Calibration of the magnetic field and measurements of magnetic field spatial uniformity are performed using a Texas Instruments TMAG5273A1 3-axis Hall-effect sensor.

Oscillating magnetic fields along the *x* direction are generated using a bridged loop-gap resonator (BLGR). The resonator used for the results in Fig. 2 is constructed from a 32 mm length copper tube with an inner (outer) diameter of 31 mm (37 mm). A single ∼ 0.6 mm wide gap is cut along the tube’s length and filled with polytetrafluoroethylene (PTFE) dielectric. A curved copper bridge, 10 mm wide and 30 mm long, is positioned symmetrically over the gap on the outside of the tube, with PTFE dielectric separating the bridge from the tube. Resonators with modified geometries were used to achieve different resonance frequencies for the data shown in Fig. 3 **A**. We note that the BLGR is very effective at heating the air within the RF shield. To prevent undesirable sample heating, the RF shield is continuously flushed with filtered dry air.

Fiber-coupled multimode laser diodes at 440 nm (Wavespectrum RLS/445NM-3500MW) and 520 nm (Wavespectrum RLS/520NM-800MW) generate excitation light. We note that an additional 561 nm source (Coherent Sapphire) was used for *C. elegans* experiments where mentioned. The lasers are combined using a multimode fiber combiner (Thorlabs MP3LF1) to ensure coaxial propagation of all wavelengths onto the sample. A dichroic beamsplitter (Semrock FF593-Di03-25x36 for experiments with RFPs and MagLOV-mScarlet fusion or Thorlabs DMLP505 for MagLOV) reflects the laser light and transmits epi-fluorescence. For experiments involving purified RFPs and *E. coli* colonies, fluorescence is collected using a zoom lens (Thorlabs MVL7002) and imaged onto a FLIR BFS-U3-32S4M-C camera. For experiments with purified RFPs in Figs. 2 **G - I** and for *C. elegans* experiments, fluorescence is collected using a 0.1 NA stereo microscope objective lens placed between the sample and the dichroic. Images are recorded using a pco.edge 5.5 sCMOS camera.

### 1.2 RF electronics

RF signals are initially generated using an HP E4421B synthesizer. The synthesizer output is amplified using a Mini-Circuits ZHL-100W-GaN+ amplifier. For transient experiments an RF switch (Mini-Circuits ZYSW-2-50DR) placed before the amplifier enables 5 ns switching of the RF using TTL logic signals. Back reflections of RF to the amplifier are limited by placing an RF circulator after the amplifier. After the circulator, a dual directional coupler (HP 778D) samples both the forward and reflected power from the resonator (using Mini-Circuits ZX47-50LN+ power detectors). These measurements were used to calculate the power dissipated in the resonator. The direct output port of the coupler is connected to a coaxial cable terminated in a ∼ 25 mm inner diameter coupling loop positioned below the BLGR. Optimization of the coupling is achieved by varying the distance between the loop and the resonator.

The quality factor (*Q*) of the BLGR resonator was determined by measuring the duration *τ* for the RF voltage amplitude across the resonator coupling loop to reduce by a factor of 1/*e* after switching off the RF. The quality factor can then be calculated using *Q* = 2*π f*_RF_*τ*/2. The RF magnetic field amplitude at the sample position *B*_1_ is related to the dissipated power (*P*) by 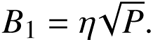 Using the method of perturbing spheres (*33, 34*), we measure 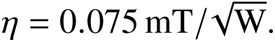

### 1.3 Data analysis and fitting procedures

Experimental data showing *B*_0_ dependent fluorescence are recorded while a sequence of sawtooth current ramps are applied to the Helmholtz coils to generate a time-dependent magnetic field profile *B*_0_(*t*).

Effects of laser-induced fluorophore bleaching are partially removed by subtracting a function F_*b*_ (*B*_0_(*t*)). For in vitro experiments this function is a straight line fitted between data points where *B*_0_ = 0 at either ends of each current ramp sequence. For experiments with *C. elegans* and for transient-response experiments (Figs. **G** - **I**), a function consisting of the sum of an exponential decay and a third-order polynomial is used to characterize the bleaching for two consecutive current ramps sequences.

After subtraction of the bleaching function data sets are analyzed by nonlinear least squares fitting (Scipy.optimize.curve fit) using the function

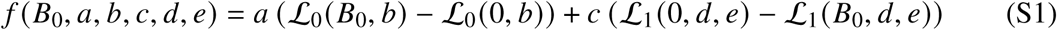

where

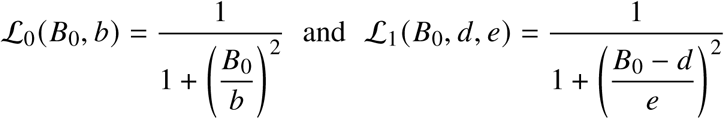

are Lorentzian functions that empirically model the MFE and the magnetic resonance respectively. The fitting algorithm returns a set of optimal fit parameters **p**^∗^ = {*a*^∗^, *b*^∗^, *c*^∗^, *d*^∗^, *e*^∗^} and a covariance matrix **cov**. Properties of the MFE and the resonance (Figs 2 **B-E** and 3**A**) are related to the fit parameters by *B*_1/2_ = *b*^∗^, *B*_0res_ = *d*^∗^, and FWHM = 2*e*^∗^. The MFE is the fraction change in fluorescence due to the magnetic field

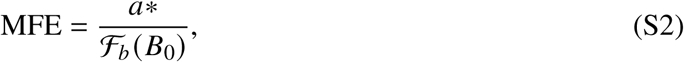

and the RYDMR amplitude is given by

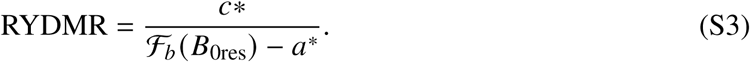

where the denominator is the estimate of the fluorescence with only the static field. Standard errors for the fit parameters are obtained from the square root of the diagonal elements of the covariance matrix 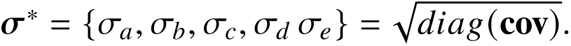 Uncertainties associated with parameters derived from the optimal fit parameters are calculated using error propagation formulae (*35*).

The fitted values of the MFE in Fig. 2 **D** are extrapolations as *B*_0_ cannot be increased beyond about 30 mT in our setup. However, the fitted values of the MFE shown in Fig. 2 **D** are consistent with measurements of the MFE performed using a ∼ 100 mT magnetic field with a solution of mScarlet-I and FMN (see S 1.5.2).

Transient-response experiments demonstrate that there is a delayed response in fluorescence intensity to a step change of the RF field (Fig. 2 **G**). This effect was incorporated in the fitting procedure by applying a first-order, low-pass filter with time constant *τ* to Eq. S1 before each evaluation of the objective function in the fitting algorithm. The values of *τ* obtained from the fit are shown in Fig. 2 **H**. The low-pass filtering effect results in a slight reduction in the measured value of *B*_0res_ (∼ 2 % for *τ* ∼ 1.5 s) and in the measured RYDMR amplitude. Scan periods for *B*_0_(*t*) were selected to be ∼ 1 min - significantly longer than the *τ* = 1.2 s required for the fluorescence to reach as new steady state after a step change in the RF.

### 1.4 Red Fluorescent Proteins

#### 1.4.1 Materials, plasmids and protein sequences

Flavin Mononucleotide (FMN) was purchased from Aaron Chemicals (catalog number: AR00AEZ8). All RFP plasmids were based on the pBAD backbone, except for mScarlet3, which was constructed on the pET-28a(+) plasmid. To avoid repetition, the complete plasmid sequences for mScarlet and mScarlet3 are detailed below, while only the protein sequences are provided for all other RFPs used in this study.

##### Whole plasmid sequence for mScarlet in pBAD backbone

gccgacatcaccgatggggaagatcgggctcgccacttcgggctcatgagcgcttgtttcggcgtgggtatggtggcagg

ccccgtggccgggggactgttgggcgccatctccttctgcctcgcgcgtttcggtgatgacggtgaaaacctctgacaca

tgcagctcccggagacggtcacagcttgtctgtaagcggatgccgggagcagacaagcccgtcagggcgcgtcagcgggt

gttggcgggtgtcggggcgcagccatgacccagtcacgtagcgatagcggagtgtatactggcttaactatgcggcatca

gagcagattgtactgagagtgcaccagatgcggtgtgaaataccgcacagatgcgtaaggagaaaataccgcatcaggcg

ctcttccgcttcctcgctcactgactcgctgcgctcggtcgttcggctgcggcgagcggtatcagctcactcaaaggcgg

taatacggttatccacagaatcaggggataacgcaggaaagaacatgtgagcaaaaggccagcaaaaggccaggaaccgt

aaaaaggccgcgttgctggcgtttttccataggctccgcccccctgacgagcatcacaaaaatcgacgctcaagtcagag

gtggcgaaacccgacaggactataaagataccaggcgtttccccctggaagctccctcgtgcgctctcctgttccgaccc

tgccgcttaccggatacctgtccgcctttctcccttcgggaagcgtggcgctttctcatagctcacgctgtaggtatctc

agttcggtgtaggtcgttcgctccaagctgggctgtgtgcacgaaccccccgttcagcccgaccgctgcgccttatccgg

taactatcgtcttgagtccaacccggtaagacacgacttatcgccactggcagcagccactggtaacaggattagcagag

cgaggtatgtaggcggtgctacagagttcttgaagtggtggcctaactacggctacactagaaggacagtatttggtatc

tgcgctctgctgaagccagttaccttcggaaaaagagttggtagctcttgatccggcaaacaaaccaccgctggtagcgg

tggtttttttgtttgcaagcagcagattacgcgcagaaaaaaaggatctcaagaagatcctttgatcttttctacggggt

ctgacgctcagtggaacgaaaactcacgttaagggattttggtcatgagattatcaaaaaggatcttcacctagatcctt

ttaaattgtaaacgttaatattttgttaaaattcgcgttaaatttttgttaaatcagctcattttttaaccaataggccg

aaatcggcaaaatcccttataaatcaaaagaatagcccgagatagggttgagtgttgttccagtttggaacaagagtcca

ctattaaagaacgtggactccaacgtcaaagggcgaaaaaccgtctatcagggcgatggcccactacgtgaaccatcacc

caaatcaagttttttggggtcgaggtgccgtaaagcactaaatcggaaccctaaagggagcccccgatttagagcttgac

ggggaaagccggcgaacgtggcgagaaaggaagggaagaaagcgaaaggagcgggcgctagggcgctggcaagtgtagcg

gtcacgctgcgcgtaaccaccacacccgccgcgcttaatgcgccgctacagggcgcgtaaatcaatctaaagtatatatg

agtaaacttggtctgacagttaccaatgcttaatcagtgaggcacctatctcagcgatctgtctatttcgttcatccata

gttgcctgactccccgtcgtgtagataactacgatacgggagggcttaccatctggccccagtgctgcaatgataccgcg

agacccacgctcaccggctccagatttatcagcaataaaccagccagccggaagggccgagcgcagaagtggtcctgcaa

ctttatccgcctccatccagtctattaattgttgccgggaagctagagtaagtagttcgccagttaatagtttgcgcaac

gttgttgccattgctgcaggcatcgtggtgtcacgctcgtcgtttggtatggcttcattcagctccggttcccaacgatc

aaggcgagttacatgatcccccatgttgtgcaaaaaagcggttagctccttcggtcctccgatcgttgtcagaagtaagt

tggccgcagtgttatcactcatggttatggcagcactgcataattctcttactgtcatgccatccgtaagatgcttttct

gtgactggtgagtactcaaccaagtcattctgagaatagtgtatgcggcgaccgagttgctcttgcccggcgtcaacacg

ggataataccgcgccacatagcagaactttaaaagtgctcatcattggaaaacgttcttcggggcgaaaactctcaagga

tcttaccgctgttgagatccagttcgatgtaacccactcgtgcacccaactgatcttcagcatcttttactttcaccagc

gtttctgggtgagcaaaaacaggaaggcaaaatgccgcaaaaaagggaataagggcgacacggaaatgttgaatactcat

actcttcctttttcaatattattgaagcatttatcagggttattgtctcatgagcggatacatatttgaatgtatttaga

aaaataaacaaaagagtttgtagaaacgcaaaaaggccatccgtcaggatggccttctgcttaatttgatgcctggcagt

ttatggcgggcgtcctgcccgccaccctccgggccgttgcttcgcaacgttcaaatccgctcccggcggatttgtcctac

tcaggagagcgttcaccgacaaacaacagataaaacgaaaggcccagtctttcgactgagcctttcgttttatttgatgc

ctggcagttccctactctcgcatggggagaccccacactaccatcggcgctacggcgtttcacttctgagttcggcatgg

ggtcaggtgggaccaccgcgctactgccgccaggcaaattctgttttatcagaccgcttctgcgttctgatttaatctgt

atcaggctgaaaatcttctctcatccgccaaaacagccaagcttcgaattcttacttgtacagctcgtccatgccgccgg

tggagtggcggccctcggagcgttcgtactgttccaccacggtgtagtcctcgttgtgggaggtgatgtccaacttgcgg

tcgacgttgtaggcgccgggcatctgcacgggcttcttggccttgtaggtggtcttgaagtccgccaggtagcggccgcc

gtccttcaggcgcagggccatcttaatgtcgcccttcagcacgccgtcctcggggtacaaccgctcggtggacgcttccc

agcccattgtcttcttctgcattacggggccgtcaggagggaagttggtgccgcggagcttcaccttgtagatcagggtg

ccgtcctccagggaggtgtcctgggtcacggtcacggcgccgccgtcctcgaagttcatcacgcgctcccacttgaagcc

ctcggggaaggactgcttatagtagtcggggatgtcggcggggtgcttggtgaaggccctggagccgtacatgaactgag

gggacaggatgtcccaggagaagggcagggggccacccttggtcaccttcagcttggcggtctgggtgccctcgtagggg

cggccctcgccctcgccctcgatctcgaactcgtggccgttcatggagccctccatgtgcaccttgaaccgcatgaactc

cttgatcactgcctcgcccttgctcaccatttttttgggatccttatcgtcatcgtcgtacagatcccgacccatttgct

gtccaccagtcatgctagccataccatgatgatgatgatgatgagaaccccgcatatgtatatctccttcttaaagttaa

acaaaattatttctagcccaaaaaaacgggtatggagaaacagtagagagttgcgataaaaagcgtcaggtagtatccgc

taatcttatggataaaaatgctatggcatagcaaagtgtgacgccgtgcaaataatcaatgtggacttttctgccgtgat

tatagacacttttgttacgcgtttttgtcatggctttggtcccgctttgttacagaatgcttttaataagcggggttacc

ggtttggttagcgagaagagccagtaaaagacgcagtgacggcaatgtctgatgcaatatggacaattggtttcttctct

gaatggcgggagtatgaaaagtatggctgaagcgcaaaatgatcccctgctgccgggatactcgtttaatgcccatctgg

tggcgggtttaacgccgattgaggccaacggttatctcgatttttttatcgaccgaccgctgggaatgaaaggttatatt

ctcaatctcaccattcgcggtcagggggtggtgaaaaatcagggacgagaatttgtttgccgaccgggtgatattttgct

gttcccgccaggagagattcatcactacggtcgtcatccggaggctcgcgaatggtatcaccagtgggtttactttcgtc

cgcgcgcctactggcatgaatggcttaactggccgtcaatatttgccaatacggggttctttcgcccggatgaagcgcac

cagccgcatttcagcgacctgtttgggcaaatcattaacgccgggcaaggggaagggcgctattcggagctgctggcgat

aaatctgcttgagcaattgttactgcggcgcatggaagcgattaacgagtcgctccatccaccgatggataatcgggtac

gcgaggcttgtcagtacatcagcgatcacctggcagacagcaattttgatatcgccagcgtcgcacagcatgtttgcttg

tcgccgtcgcgtctgtcacatcttttccgccagcagttagggattagcgtcttaagctggcgcgaggaccaacgtatcag

ccaggcgaagctgcttttgagcaccacccggatgcctatcgccaccgtcggtcgcaatgttggttttgacgatcaactct

atttctcgcgggtatttaaaaaatgcaccggggccagcccgagcgagttccgtgccggttgtgaagaaaaagtgaatgat

gtagccgtcaagttgtcataattggtaacgaatcagacaattgacggcttgacggagtagcatagggtttgcagaatccc

tgcttcgtccatttgacaggcacattatgcatcgatgataagctgtcaaacatgagcagatcctctacgccggacgcatc

gtggccggcatcaccggcgccacaggtgcggttgctggcgcctatatc

##### mScarlet Protein Sequence

MRGSHHHHHH GMASMTGGQQ MGRDLYDDDD KDPKKMVSKG EAVIKEFMRF KVHMEGSMNG

HEFEIEGEGE GRPYEGTQTA KLKVTKGGPL PFSWDILSPQ FMYGSRAFTK HPADIPDYYK

QSFPEGFKWE RVMNFEDGGA VTVTQDTSLE DGTLIYKVKL RGTNFPPDGP VMQKKTMGWE

ASTERLYPED GVLKGDIKMA LRLKDGGRYL ADFKTTYKAK KPVQMPGAYN VDRKLDITSH

NEDYTVVEQY ERSEGRHSTG GMDELYK

##### mScarlet-I Protein Sequence

MRGSHHHHHH GMASMTGGQQ MGRDLYDDDD KDPKKMVSKG EAVIKEFMRF KVHMEGSMNG

HEFEIEGEGE GRPYEGTQTA KLKVTKGGPL PFSWDILSPQ FMYGSRAFIK HPADIPDYYK

QSFPEGFKWE RVMNFEDGGA VTVTQDTSLE DGTLIYKVKL RGTNFPPDGP VMQKKTMGWE

ASTERLYPED GVLKGDIKMA LRLKDGGRYL ADFKTTYKAK KPVQMPGAYN VDRKLDITSH

NEDYTVVEQY ERSEGRHSTG GMDELYK

##### mCherry Protein Sequence

MRGSHHHHHH GMASMTGGQQ MGRDLYDDDD KDPMVSKGEE DNMAIIKEFM RFKVHMEGSV

NGHEFEIEGE GEGRPYEGTQ TAKLKVTKGG PLPFAWDILS PQFMYGSKAY VKHPADIPDY

LKLSFPEGFK WERVMNFEDG GVVTVTQDSS LQDGEFIYKV KLRGTNFPSD GPVMQKKTMG

WEASSERMYP EDGALKGEIK QRLKLKDGGH YDAEVKTTYK AKKPVQLPGA YNVNIKLDIT

SHNEDYTIVE QYERAEGRHS TGGMDELYK

##### mCherry-XL Protein Sequence

MRGSHHHHHH GMASMTGGQQ MGRDLYDDDD KDPKKMVSKG EEDNMAIIKE FMRFKVHMEG

SVNGHEFEIE GEGEGRPYEG TQTAKLKVTK GGPLPFAWDI LSPQFMYGSK AYVKHPADIP

DYLKLSFPEG FKWERVMNFE DGGVVTVTQD SSLQDGEFIY KVKLKGTNFP SDGPVMQKKT

MGSEASSERM YPEDGALKGE VKYRLKLKDG GHYDAEVKTT YKAKKPVQLP GAYNVNRKLD

ITSHNEDYTI VEQYERAEGR HSTGGMDELY K

##### mCherry-D Protein Sequence

MRGSHHHHHH GMASMTGGQQ MGRDLYDDDD KDPKKMVSKA EEDNMAIIKE FMRFKTRMEG

SVNGHEFEIE GEGEGRPYEG TQTAKLKVTK GGPLPFAWDI LSPQFMYGSR AYVKHPADIP

DYLKLSFPEG FKWERVMKSE DGGVVTVTQD SSLQDGEFIY KVKLRGTNFP SDGPVMQKKT

MGWEASSERM YPEDGALKGE MKMRLRLKDG GHYDWEVKTT YKAKKPVQLP GAYNVNRKLD

ITSHNEDYTI VEQYERAEGR HSTGGMDELY K

##### Whole plasmid sequence for mScarlet3 in pET-28a(+) backbone

Tggcgaatgggacgcgccctgtagcggcgcattaagcgcggcgggtgtggtggttacgcgcagcgtgaccgctacacttg

ccagcgccctagcgcccgctcctttcgctttcttcccttcctttctcgccacgttcgccggctttccccgtcaagctcta

aatcgggggctccctttagggttccgatttagtgctttacggcacctcgaccccaaaaaacttgattagggtgatggttc

acgtagtgggccatcgccctgatagacggtttttcgccctttgacgttggagtccacgttctttaatagtggactcttgt

tccaaactggaacaacactcaaccctatctcggtctattcttttgatttataagggattttgccgatttcggcctattgg

ttaaaaaatgagctgatttaacaaaaatttaacgcgaattttaacaaaatattaacgtttacaatttcaggtggcacttt

tcggggaaatgtgcgcggaacccctatttgtttatttttctaaatacattcaaatatgtatccgctcatgaattaattct

tagaaaaactcatcgagcatcaaatgaaactgcaatttattcatatcaggattatcaataccatatttttgaaaaagccg

tttctgtaatgaaggagaaaactcaccgaggcagttccataggatggcaagatcctggtatcggtctgcgattccgactc

gtccaacatcaatacaacctattaatttcccctcgtcaaaaataaggttatcaagtgagaaatcaccatgagtgacgact

gaatccggtgagaatggcaaaagtttatgcatttctttccagacttgttcaacaggccagccattacgctcgtcatcaaa

atcactcgcatcaaccaaaccgttattcattcgtgattgcgcctgagcgagacgaaatacgcgatcgctgttaaaaggac

aattacaaacaggaatcgaatgcaaccggcgcaggaacactgccagcgcatcaacaatattttcacctgaatcaggatat

tcttctaatacctggaatgctgttttcccggggatcgcagtggtgagtaaccatgcatcatcaggagtacggataaaatg

cttgatggtcggaagaggcataaattccgtcagccagtttagtctgaccatctcatctgtaacatcattggcaacgctac

ctttgccatgtttcagaaacaactctggcgcatcgggcttcccatacaatcgatagattgtcgcacctgattgcccgaca

ttatcgcgagcccatttatacccatataaatcagcatccatgttggaatttaatcgcggcctagagcaagacgtttcccg

ttgaatatggctcataacaccccttgtattactgtttatgtaagcagacagttttattgttcatgaccaaaatcccttaa

cgtgagttttcgttccactgagcgtcagaccccgtagaaaagatcaaaggatcttcttgagatcctttttttctgcgcgt

aatctgctgcttgcaaacaaaaaaaccaccgctaccagcggtggtttgtttgccggatcaagagctaccaactctttttc

cgaaggtaactggcttcagcagagcgcagataccaaatactgtccttctagtgtagccgtagttaggccaccacttcaag

aactctgtagcaccgcctacatacctcgctctgctaatcctgttaccagtggctgctgccagtggcgataagtcgtgtct

taccgggttggactcaagacgatagttaccggataaggcgcagcggtcgggctgaacggggggttcgtgcacacagccca

gcttggagcgaacgacctacaccgaactgagatacctacagcgtgagctatgagaaagcgccacgcttcccgaagggaga

aaggcggacaggtatccggtaagcggcagggtcggaacaggagagcgcacgagggagcttccagggggaaacgcctggta

tctttatagtcctgtcgggtttcgccacctctgacttgagcgtcgatttttgtgatgctcgtcaggggggcggagcctat

ggaaaaacgccagcaacgcggcctttttacggttcctggccttttgctggccttttgctcacatgttctttcctgcgtta

tcccctgattctgtggataaccgtattaccgcctttgagtgagctgataccgctcgccgcagccgaacgaccgagcgcag

cgagtcagtgagcgaggaagcggaagagcgcctgatgcggtattttctccttacgcatctgtgcggtatttcacaccgca

tatatggtgcactctcagtacaatctgctctgatgccgcatagttaagccagtatacactccgctatcgctacgtgactg

ggtcatggctgcgccccgacacccgccaacacccgctgacgcgccctgacgggcttgtctgctcccggcatccgcttaca

gacaagctgtgaccgtctccgggagctgcatgtgtcagaggttttcaccgtcatcaccgaaacgcgcgaggcagctgcgg

taaagctcatcagcgtggtcgtgaagcgattcacagatgtctgcctgttcatccgcgtccagctcgttgagtttctccag

aagcgttaatgtctggcttctgataaagcgggccatgttaagggcggttttttcctgtttggtcactgatgcctccgtgt

aagggggatttctgttcatgggggtaatgataccgatgaaacgagagaggatgctcacgatacgggttactgatgatgaa

catgcccggttactggaacgttgtgagggtaaacaactggcggtatggatgcggcgggaccagagaaaaatcactcaggg

tcaatgccagcgcttcgttaatacagatgtaggtgttccacagggtagccagcagcatcctgcgatgcagatccggaaca

taatggtgcagggcgctgacttccgcgtttccagactttacgaaacacggaaaccgaagaccattcatgttgttgctcag

gtcgcagacgttttgcagcagcagtcgcttcacgttcgctcgcgtatcggtgattcattctgctaaccagtaaggcaacc

ccgccagcctagccgggtcctcaacgacaggagcacgatcatgcgcacccgtggggccgccatgccggcgataatggcct

gcttctcgccgaaacgtttggtggcgggaccagtgacgaaggcttgagcgagggcgtgcaagattccgaataccgcaagc

gacaggccgatcatcgtcgcgctccagcgaaagcggtcctcgccgaaaatgacccagagcgctgccggcacctgtcctac

gagttgcatgataaagaagacagtcataagtgcggcgacgatagtcatgccccgcgcccaccggaaggagctgactgggt

tgaaggctctcaagggcatcggtcgagatcccggtgcctaatgagtgagctaacttacattaattgcgttgcgctcactg

cccgctttccagtcgggaaacctgtcgtgccagctgcattaatgaatcggccaacgcgcggggagaggcggtttgcgtat

tgggcgccagggtggtttttcttttcaccagtgagacgggcaacagctgattgcccttcaccgcctggccctgagagagt

tgcagcaagcggtccacgctggtttgccccagcaggcgaaaatcctgtttgatggtggttaacggcgggatataacatga

gctgtcttcggtatcgtcgtatcccactaccgagatatccgcaccaacgcgcagcccggactcggtaatggcgcgcattg

cgcccagcgccatctgatcgttggcaaccagcatcgcagtgggaacgatgccctcattcagcatttgcatggtttgttga

aaaccggacatggcactccagtcgccttcccgttccgctatcggctgaatttgattgcgagtgagatatttatgccagcc

agccagacgcagacgcgccgagacagaacttaatgggcccgctaacagcgcgatttgctggtgacccaatgcgaccagat

gctccacgcccagtcgcgtaccgtcttcatgggagaaaataatactgttgatgggtgtctggtcagagacatcaagaaat

aacgccggaacattagtgcaggcagcttccacagcaatggcatcctggtcatccagcggatagttaatgatcagcccact

gacgcgttgcgcgagaagattgtgcaccgccgctttacaggcttcgacgccgcttcgttctaccatcgacaccaccacgc

tggcacccagttgatcggcgcgagatttaatcgccgcgacaatttgcgacggcgcgtgcagggccagactggaggtggca

acgccaatcagcaacgactgtttgcccgccagttgttgtgccacgcggttgggaatgtaattcagctccgccatcgccgc

ttccactttttcccgcgttttcgcagaaacgtggctggcctggttcaccacgcgggaaacggtctgataagagacaccgg

catactctgcgacatcgtataacgttactggtttcacattcaccaccctgaattgactctcttccgggcgctatcatgcc

ataccgcgaaaggttttgcgccattcgatggtgtccgggatctcgacgctctcccttatgcgactcctgcattaggaagc

agcccagtagtaggttgaggccgttgagcaccgccgccgcaaggaatggtgcatgcaaggagatggcgcccaacagtccc

ccggccacggggcctgccaccatacccacgccgaaacaagcgctcatgagcccgaagtggcgagcccgatcttccccatc

ggtgatgtcggcgatataggcgccagcaaccgcacctgtggcgccggtgatgccggccacgatgcgtccggcgtagagga

tcgagatctcgatcccgcgaaattaatacgactcactataggggaattgtgagcggataacaattcccctctagaaataa

ttttgtttaactttaagaaggagatataccatgaggggatcacatcaccaccatcaccactctagcggtttggttccgcg

tgcggttattaaagagttcatgcgttttaaggtgcacatggaagggtctatgaatggtcatgaattcgagatcgaaggcg

agggcgagggtcgtccgtacgaaggcacccagaccgcgaagctgcgcgttaccaaaggtggtccgctgccgtttagctgg

gatattctgtccccgcaatttatgtatggtagccgtgccttcaccaagcacccggcggacatcccggactactggaaaca

atcgttcccggaaggtttcaagtgggagcgcgtgatgaactttgaggacggcggcgcggtgagcgttgcgcaggatacct

ccttggaagacggcactctgatttataaagtcaaattgcgcggtacgaacttccctccggatggcccagtaatgcagaaa

aagacgatgggttgggaggctagcaccgaacgtttatatccggaggacgtcgttctgaagggtgatatcaaaatggcact

gagacttaaggacggcggtcgctacctggcagattttaagaccacgtaccgcgctaaaaagccggtgcagatgccgggtg

cgttcaacattgatcgtaaactggatatcacctcccacaatgaagactacaccgttgtggaacaatatgagcgtagcgtg

gcccgtcatagctaacaaagcccgaaaggaagctgagttggctgctgccaccgctgagcaataactagcataaccccttg

gggcctctaaacgggtcttgaggggttttttgctgaaaggaggaactatatccggat

##### mScarlet3 Protein Sequence

MRGSHHHHHH SSGLVPRAVI KEFMRFKVHM EGSMNGHEFE IEGEGEGRPY EGTQTAKLRV

TKGGPLPFSW DILSPQFMYG SRAFTKHPAD IPDYWKQSFP EGFKWERVMN FEDGGAVSVA

QDTSLEDGTL IYKVKLRGTN FPPDGPVMQK KTMGWEASTE RLYPEDVVLK GDIKMALRLK

DGGRYLADFK TTYRAKKPVQ MPGAFNIDRK LDITSHNEDY TVVEQYERSV ARHS

#### 1.4.2 Protein expression and purification

Each RFP plasmid was transformed into *Escherichia coli* BL21(DE3) cells (Invitrogen). Selected colonies were inoculated into two 5 mL cultures of Luria-Bertani (LB) medium supplemented with 100 µg/mL ampicillin and incubated overnight at 37 °C with shaking at 225 rpm. The following day, the overnight cultures were combined and transferred into 1 L of LB medium, supplemented with ampicillin 100 µg/ml. The culture was grown at 37 °C and 220 rpm until the optical density at 600 nm (OD600) reached 0.6. At this point, arabinose was added to a final concentration of 2 mg/ml to induce protein expression. The cells were then incubated overnight at 18 °C with shaking at 220 rpm.

On the next day, the cells were harvested by centrifugation at 4000 rpm for 30 minutes and stored at -80 °C. For lysis, cells were resuspended in lysis buffer (0.1 M Tris, pH 8.0) and disrupted using a homogenizer set to a pressure of 20 MPa with a flow rate of 35 mL/min. Five passes through the homogenizer were used to ensure complete cell lysis. The lysate was centrifuged at 16,000 rpm for 45 minutes to pellet cellular debris. The supernatant was filtered through a 0.22 µm membrane (Millipore Corp.) and loaded onto a His-tag affinity column pre-equilibrated with lysis buffer. Proteins without His-tag were removed by washing the column with 5 column volumes of wash buffer (0.1 M Tris, 20 mM imidazole, pH 8.0). The target protein was eluted using 3 column volumes of elution buffer (0.1 M Tris, 300 mM imidazole, pH 8.0). Finally, the protein was buffer-exchanged into 0.1 M Tris buffer, pH 8.0.

#### 1.4.3 MFEs and RYDMR in other RFPs

We have measured the MFEs from the emission of various RFPs together with FMN. We note that the magnitude of MFE is influenced by several factors including the mutations of the fluorescent protein, the FMN to protein concentration ratio, temperature, and green and blue optical excitation intensities. Among the RFPs tested, the largest MFEs (measured in the regime where Δ*g* mechanism is insignificant) have been observed in mScarlet and its variants mScarlet-I, mScarlet3, reaching approximately 20 %, while mCherry exhibits the lowest MFE saturating at around 1.5%. MFEs and RYDMR data for mCherry and two variants: mCherry-XL, and mCherry-D (*36, 37*) are shown in Fig. S2. The mutations in these mCherry variants resulted in increases in both the MFE and the RYDMR amplitude compared to mCherry, indicating that mutations can influence the magnetic response of fluorescent proteins. No obvious essential amino acids leading to these differences have been identified to date. To avoid RF heating, all in vitro RYDMR experiments are conducted by dissolving RFP and FMN in distilled water.

### 1.5 Proposed mechanism for MFEs in RFP-flavin systems

A hypothesized spin-correlated radical pair (RP)-based reaction scheme in the FMN-RFP system is illustrated in Fig. S3. In the investigated RFP-flavin systems, photosensitization with blue light, presumably to convert FMN to an FMN photoproduct, seems to be necessary to observe MFEs. While RFPs can weakly absorb blue light and fluoresce, the largest MFEs in their emission are observed using green light excitation, following or together with blue light excitation. This suggests RP formation between the RFP and FMN photoproduct during the green light excitation period. Furthermore, after the system has been pre-excited using blue light, introducing a period with no optical excitation ranging from seconds (Fig. S6) to several minutes before green light excitation, still results in MFEs in fluorescence. This suggests that the FMN photoproduct is likely not an FMN excited state. The precise chemical identity of the photoproduct remains unknown, and we tentatively call it flavinX.

Application of green light to a mixture of RFP and flavinX can result in excitation of the RFP chromophore, which we denote RFP^*c*^, to form RFP^*c*∗^ which can fluoresce and return to the RFP^*c*^ ground state. We distinguish the protein barrel, denoted RFP, from the RFP chromophore, RFP^*c*^, in the following. Additionally, we hypothesize that green light excitation can lead to the formation of a RP between flavinX, initially in the triplet state ^3^flavinX, and RFP, as suggested by CIDNP studies of amino acid:flavin systems, where flavin or its photoproduct is formed in the triplet state (*29, 30*). This process likely involves electron transfer or hydrogen atom abstraction from an amino acid on the surface of the RFP by ^3^flavinX. We invoke the triplet state of flavinX as this is a bimolecular reaction and the likely lifetime of the singlet state of flavinX is too short for RP formation to be efficient. We note that it is possible that the initial spin state of the radical pair is a singlet state, but the same RP would be involved and MFEs would still be expected.

Hyperfine interactions drive coherent interconversion between the singlet state (*S*) and triplet state (*T*) of the RP, which can be sensitive to external magnetic fields. From *S*, the RP can rapidly decay —on nanosecond to microsecond timescales—resulting in repopulation of the RFP ground state. From *T*, the RP can either convert to *S* or decay, resulting in a dark species whose chemical identity remains unknown. We hypothesize that this dark species could transition back to the RFP ground state over longer timescales (presumably on the millisecond to second timescales), as ∼10s of cycles of magnetically-induced fluorescence modulation over a total period of several minutes do not lead to a significant decrease in fluorescence intensity or measured MFE for each cycle. The precise feature(s) of the RFP component of the radical pair is not known at this time. This could be a radical on the surface of the protein that affects the excited state properties of RFP^*c*^ or it could directly involve the RFP chromophore, e.g. by secondary electron transfer.

While still preliminary, aspects of this model are supported by in vitro absorption and fluorescence measurements performed on solutions of RFP with FMN. These measurements are described in detail in the subsections below and the key observations are summarized here. During the blue light excitation period, fluorescence measurements (Fig. S6 B) show a reduction in the concentration of ground state oxidized FMN (fluorescence maximum near 530 nm), indicating the formation of a photoproduct. With the blue light source switched off, applying green light results in an initial decrease in mScarlet fluorescence (Fig. S6 B) and a decrease in absorption (Fig. S5 A) associated with the fluorescent protein (around 569 nm). This reduction in absorption is inversely correlated with the formation of a new absorption peak centered near 520 nm (Fig. S5 A). When an external magnetic field is applied, RFP fluorescence and absorption (around 569 nm) both decrease (up to ∼20 %), while the absorption feature at 520 nm increases. We hypothesize that the 520 nm feature corresponds to the dark species mentioned above. Spectrally resolved fluorescence measurements show no obvious spectral shifts in the mScarlet emission resulting from the magnetic field (Fig. S6 C). These observations suggest that following FMN photoproduct formation, the magnetic field serves to control the population of RFP molecules that can be excited and fluoresce in the presence of green light.

#### 1.5.1 Absorption measurements

Steady-state absorption spectra (obtained using a Perkin Elmer LAMBDA 365+ UV/Vis Spectrometer) for a mixture of mScarlet and FMN, as well as for the individual components (FMN and mScarlet), are shown in Fig. S4. indicating no significant alteration in their absorption properties due to potential interactions.

Time-resolved absorption spectra of the mScarlet and FMN mixture are presented in Fig. S5. Spectra were recorded following 30 s pre-excitation with a 440 nm laser with intensity at the sample of 9 W/cm^2^. Subsequently, probe light from a 3500 K white LED was used to irradiate the sample. Simultaneously with the probe, light from a 520 nm laser with intensity 7 W/cm^2^ was incident on the sample. This source was pulsed with a frequency of 10 Hz and a 50 % duty cycle. Absorption of the white light probe was recorded only during the intervals where the 520 nm laser was off using an OceanFX spectrometer (Ocean Optics). A magnetic field, generated using Helmholtz coils, was switched between 0 and ∼ 40 mT with 50 % duty cycle and a 40 s period to enable measurement of magnetic field effects throughout the experiment.

Immediately following the initialization of 520 nm light exposure, there is an absorption maximum at 569 nm, corresponding to the mScarlet absorption peak (Fig.S5 A). As the duration of 520 nm exposure increases, the absorption peak at 569 nm falls almost 50 % after 0.7 s and a new absorption peak is formed near 520 nm (Fig.S5 A). The increase in absorption at 520 nm is commensurate with the decrease in absorption at 569 nm suggesting the formation of a non-fluorescent version of the FP, inversely correlated with the RFP ground state population measured at 569 nm. Furthermore, the change in absorption at 520 and 569 nm induced by the magnetic field are also anticorrelated (Fig.S5 B, C, and D) suggesting that the reduction in RFP fluorescence due to the magnetic field results from a reduction in the population of ground state RFPs that can be excited and fluoresce. Significantly weaker modulation correlated with the magnetic field switching is present at 445 nm (corresponding to the FMN absorption maximum) and at 594 nm (the mScarlet emission maximum).

Control experiments with solutions containing only FMN or mScarlet using the same pre-excitation and green light irradiation showed no evidence of increased absorption at 520 nm. To rule out potential artifacts from the 520 nm excitation laser, the experiments were repeated using a 569 nm laser instead. The longer wavelength excitation source results in no significant changes to the results shown in Fig. S5 or to their interpretation.

#### 1.5.2 Fluorescence measurements

Spectrally-resolved fluorescence from a solution of 50 *μ*M mScarlet-I and 300 *μ*M FMN was measured during a period of blue light excitation and subsequently during a period of green light excitation (Fig. S6). A magnetic field is switched between 0 and ∼100 mT every 90 s throughout the experiment by periodically moving a permanent magnet close to the sample. A 450 nm light emitting diode (LED), with an 18 nm FWHM bandwidth and a 530 nm LED with a 35 nm FWHM bandwidth were used as excitation sources. Fluorescence was measured using an Ocean SR2 spectrometer (Ocean Optics). Rapid quenching of the mScarlet-I fluorescence to ∼ 10 % of its initial value occurs within seconds of the start of the excitation period (Fig. S6 B). For the remaining ∼ 10 min of blue light exposure, the fluorescence remains suppressed and weak modulation anti-correlated with the switching of the magnetic field is visible. After a period with no excitation, applying green light results in increased fluorescence and an increased magnetic field effect (∼20 % reduction in fluorescence). Representative spectra with and without the magnetic field are given in Fig. S6 C. The magnetic field results in no obvious spectral shifts in the emission spectra.

### 1.6 Caenorhabditis elegans

#### 1.6.1 Preparation of transgenic *C. elegans* expressing mScarlet in all cells

The worm strain WBM1143 [eft-3p::3XFLAG::wrmScarlet::unc-54 3’UTR *wbmIs65] (*24*) was used for the ubiquitous expression of a worm codon optimized version of mScarlet. This strain was obtained from the from Caenorhabditis Genetics Center (CGC). *C. elegans* were grown and maintained at 20 °C on nematode growth media (NGM) plates supplemented with the OP50 strain of *E. coli* as the food source (*38*). For imaging, larval stage 4 (L4) and day one adults were mounted in 4% agarose and paralyzed with levamisole (1 mM).

#### 1.6.2 *C. elegans* RYDMR experiments

MFEs and RYDMR were measured in multiple *C. elegans* samples with *B*_1_ = 0.1 mT (Fig. S7) and *B*_1_ = 0 mT (Fig. S8). To increase fluorescence intensity and signal-to-noise ratio, in addition to the 520 nm laser used in previous measurements, we employed a 561 nm laser for green light excitation, with a total green laser intensity of ∼ 40 W/cm^2^. For each sample, fluorescence images were recorded at various values of *B*_0_. Fitting Eq. S1 with first-order low-pass filtering to the integrated fluorescence from a region of interest (ROI) in the image gives an estimate of the MFE and the RYDMR for that ROI (Fig. S7, Columns 1 and 2). Maps of the MFE and RYDMR are obtained by dividing the image into blocks and estimating the MFE and RYDMR in each one (Fig. S7, Columns 3 and 4). Figs. 4, S7, and S8 only show RYDMR estimates from blocks where the coefficient of determination *R*^2^ > 0.6 and *p*_*F*_ < 0.01. The *p*_F_ value is measured by performing an *F* test (*39*) to determine whether the addition of fit parameters modeling the RYDMR (*c*, *d* and *e* in Eq. S1) significantly improve the fit. This is done to exclude blocks where the fit performs poorly such as in regions with low fluorescence away from the nematodes. The RYDMR and MFE distributions shown in Fig. 4 and Fig. S9 are smoothed with a 1 sigma Gaussian filter (scipy.ndimage.gaussian filter). The unfiltered RYDMR distribution corresponding to Fig. 4 F is shown in Fig. S7 row 1 column 4.

#### 1.6.2 Wild-type *C. elegans* autofluorescence

Experiments with wild-type (WT) *C. elegans* were performed to determine if the autofluorescence from WT nematodes exhibited any magnetic-field sensitivity as has been observed in mammalian cells (*40*). No significant MFEs or RYDMR in the autofluorescence of WT worms were observed using the previously described experimental and analysis procedures for the mScarlet expressing nematodes. We performed additional measurements where the usual 650 nm LP fluorescence filter was replaced with a 550 nm LP filter to collect more autofluorescence and improve the measurement signal-to-noise ratio. In these measurements, 561 nm excitation was not used. We measure changes in fluorescence consistent with MFEs of up to 0.6 % in isolated locations in WT samples (Fig. S9 A). In comparison, up to 4 % MFEs were measured using the same 550 nm LP filter with an mScarlet-expressing sample, with a similar spatial distribution to measurements obtained using the 650 nm filter. These experiments clearly demonstrate that the MFEs and RYDMR measured in mScarlet-expressing *C. elegans* is largely due to the presence of the RFP.

### 1.7 MagLOV and mScarlet-MagLOV fusion proteins

#### 1.7.1 Plasmids, sequences and *E. coli* colony preparation

Both MagLOV and mScarlet-MagLOV constructs are based on the pRSET plasmids. To avoid repetition, the full plasmid sequence for MagLOV is provided, while only the protein sequence for mScarlet-MagLOV is detailed below.

##### Whole Plasmid Sequence for MagLOV in pRSET Backbone

GATCTCGATCCCGCGAAATTAATACGACTCACTATAGGGAGACCACAACGGTTTCCCTCTAGAAATAATTTTGTTTAACT

TTAAGAAGGAGATATACATATGCGGGGTTCTCATCATCATCATCATCATGGTATGGCTAGCATGCTTGCTACTACTTTAG

AACGTATAGAGAAAAACTTCGTGATCACGGACCCGAGACTACCTGACAACCCTATAATTTTTGCAAGTGACTCATTCCTT

CAGTTGACTGAGTATTCTAGGGAAGAGATTCTAGGGtggAATcCTAGATTCTTGCAAGGACCAGAAACTGACCGTGCCAC

TGTGAGGAAAATCAGGGATGCGATCGACAACCAAACCGAGGTGACAGTGCAGCTAATAAATTACACTAAATCTGGCAAGA

AGTTCTGGAACCTATTTCATgTGCAACCCATGAGAGACCAAAAAGGAGACGTACAGTACTTCATAGGGGTAaagTTGGAT

GGTACTGAGCATGTTAGAGACGCGGCAGAACGTGAAcGGGTTATGTTAATAAAAAAGACCGCTGAAAACATAatgGAAGC

GGCAAAGGAGTTGTAACTCGAGATCTGCAGCTGGTACCATGGAATTCGAAGCTTGATCCGGCTGCTAACAAAGCCCGAAAGGAAGCTGAGTTGGCTGCTGCCACCGCTGAGCAATAACTAGCATAACCCCTTGGGGCCTCTAAACGGGTCTTGAGGGGTT

TTTTGCTGAAAGGAGGAACTATATCCGGATCTGGCGTAATAGCGAAGAGGCCCGCACCGATCGCCCTTCCCAACAGTTGC

GCAGCCTGAATGGCGAATGGGACGCGCCCTGTAGCGGCGCATTAAGCGCGGCGGGTGTGGTGGTTACGCGCAGCGTGACC

GCTACACTTGCCAGCGCCCTAGCGCCCGCTCCTTTCGCTTTCTTCCCTTCCTTTCTCGCCACGTTCGCCGGCTTTCCCCG

TCAAGCTCTAAATCGGGGGCTCCCTTTAGGGTTCCGATTTAGTGCTTTACGGCACCTCGACCCCAAAAAACTTGATTAGG

GTGATGGTTCACGTAGTGGGCCATCGCCCTGATAGACGGTTTTTCGCCCTTTGACGTTGGAGTCCACGTTCTTTAATAGT

GGACTCTTGTTCCAAACTGGAACAACACTCAACCCTATCTCGGTCTATTCTTTTGATTTATAAGGGATTTTGCCGATTTC

GGCCTATTGGTTAAAAAATGAGCTGATTTAACAAAAATTTAACGCGAATTTTAACAAAATATTAACGCTTACAATTTAGG

TGGCACTTTTCGGGGAAATGTGCGCGGAACCCCTATTTGTTTATTTTTCTAAATACATTCAAATATGTATCCGCTCATGA

GACAATAACCCTGATAAATGCTTCAATAATATTGAAAAAGGAAGAGTATGAGTATTCAACATTTCCGTGTCGCCCTTATT

CCCTTTTTTGCGGCATTTTGCCTTCCTGTTTTTGCTCACCCAGAAACGCTGGTGAAAGTAAAAGATGCTGAAGATCAGTT

GGGTGCACGAGTGGGTTACATCGAACTGGATCTCAACAGCGGTAAGATCCTTGAGAGTTTTCGCCCCGAAGAACGTTTTC

CAATGATGAGCACTTTTAAAGTTCTGCTATGTGGCGCGGTATTATCCCGTATTGACGCCGGGCAAGAGCAACTCGGTCGC

CGCATACACTATTCTCAGAATGACTTGGTTGAGTACTCACCAGTCACAGAAAAGCATCTTACGGATGGCATGACAGTAAG

AGAATTATGCAGTGCTGCCATAACCATGAGTGATAACACTGCGGCCAACTTACTTCTGACAACGATCGGAGGACCGAAGG

AGCTAACCGCTTTTTTGCACAACATGGGGGATCATGTAACTCGCCTTGATCGTTGGGAACCGGAGCTGAATGAAGCCATA

CCAAACGACGAGCGTGACACCACGATGCCTGTAGCAATGGCAACAACGTTGCGCAAACTATTAACTGGCGAACTACTTAC

TCTAGCTTCCCGGCAACAATTAATAGACTGGATGGAGGCGGATAAAGTTGCAGGACCACTTCTGCGCTCGGCCCTTCCGG

CTGGCTGGTTTATTGCTGATAAATCTGGAGCCGGTGAGCGTGGGTCTCGCGGTATCATTGCAGCACTGGGGCCAGATGGT

AAGCCCTCCCGTATCGTAGTTATCTACACGACGGGGAGTCAGGCAACTATGGATGAACGAAATAGACAGATCGCTGAGAT

AGGTGCCTCACTGATTAAGCATTGGTAACTGTCAGACCAAGTTTACTCATATATACTTTAGATTGATTTAAAACTTCATT

TTTAATTTAAAAGGATCTAGGTGAAGATCCTTTTTGATAATCTCATGACCAAAATCCCTTAACGTGAGTTTTCGTTCCAC

TGAGCGTCAGACCCCGTAGAAAAGATCAAAGGATCTTCTTGAGATCCTTTTTTTCTGCGCGTAATCTGCTGCTTGCAAAC

AAAAAAACCACCGCTACCAGCGGTGGTTTGTTTGCCGGATCAAGAGCTACCAACTCTTTTTCCGAAGGTAACTGGCTTCA

GCAGAGCGCAGATACCAAATACTGTTCTTCTAGTGTAGCCGTAGTTAGGCCACCACTTCAAGAACTCTGTAGCACCGCCT

ACATACCTCGCTCTGCTAATCCTGTTACCAGTGGCTGCTGCCAGTGGCGATAAGTCGTGTCTTACCGGGTTGGACTCAAG

ACGATAGTTACCGGATAAGGCGCAGCGGTCGGGCTGAACGGGGGGTTCGTGCACACAGCCCAGCTTGGAGCGAACGACCT

ACACCGAACTGAGATACCTACAGCGTGAGCTATGAGAAAGCGCCACGCTTCCCGAAGGGAGAAAGGCGGACAGGTATCCG

GTAAGCGGCAGGGTCGGAACAGGAGAGCGCACGAGGGAGCTTCCAGGGGGAAACGCCTGGTATCTTTATAGTCCTGTCGG

GTTTCGCCACCTCTGACTTGAGCGTCGATTTTTGTGATGCTCGTCAGGGGGGCGGAGCCTATGGAAAAACGCCAGCAACGCGGCCTTTTTACGGTTCCTGGCCTTTTGCTGGCCTTTTGCTCACATGTTCTTTCCTGCGTTATCCCCTGATTCTGTGGAT

AACCGTATTACCGCCTTTGAGTGAGCTGATACCGCTCGCCGCAGCCGAACGACCGAGCGCAGCGAGTCAGTGAGCGAGGA

AGCGGAAGAGCGCCCAATACGCAAACCGCCTCTCCCCGCGCGTTGGCCGATTCATTAATGCAG

##### mScarlet-MagLOV Protein Sequence

MRGSHHHHHH GMASMVSKGE AVIKEFMRFK VHMEGSMNGH EFEIEGEGEG RPYEGTQTAK

LKVTKGGPLP FSWDILSPQF MYGSRAFTKH PADIPDYYKQ SFPEGFKWER VMNFEDGGAV

TVTQDTSLED GTLIYKVKLR GTNFPPDGPV MQKKTMGWEA STERLYPEDG VLKGDIKMAL

RLKDGGRYLA DFKTTYKAKK PVQMPGAYNV DKKLDITSHN EDYTVVEQYE RSEGRHSTGG

GATGLATTLE RIEKNFVITD PRLPDNPIIF ASDSFLQLTE YSREEILGWN PRFLQGPETD

RATVRKIRDA IDNQTEVTVQ LINYTKSGKK FWNLLHVQPM RDQKGDVQYF IGVKLDGTEH

VRDAAGRERV MLIKKTAENI MEAAKELGTM EFEA

Each plasmid was transformed into BL21(DE3) *E. coli* cells and plated onto LB agarose plates. All MFE and RYDMR measurements were performed after one day of incubation at 37^◦^C and one day of incubation at room temperature on the LB agarose plates.

#### 1.7.2 MFEs and RYDMR

MFE and RYDMR experiments for mScarlet-MagLOV and MagLOV colonied were conducted similarly to the procedures described for RFPs in the main text. For MagLOV samples, only blue excitation light is necessary, and a 500 nm longpass filter is used to capture flavin fluorescence. Representative data from single colonies expressing the mScarlet-MagLOV fusion and MagLOV are shown in Fig. S10. Our setup allows for the measurement of MFE and RYDMR from multiple *E. coli* colonies simultaneously (Fig. S10), with the potential to be used for directed evolution to identify proteins with optimized magnetic resonance and MFEs. Our results suggest that the static MFEs observed for MagLOV by Ingaramo et al. (*27*) are due to spin-correlated radical pairs.

## 2 Supplementary Figures

**Figure S1:**
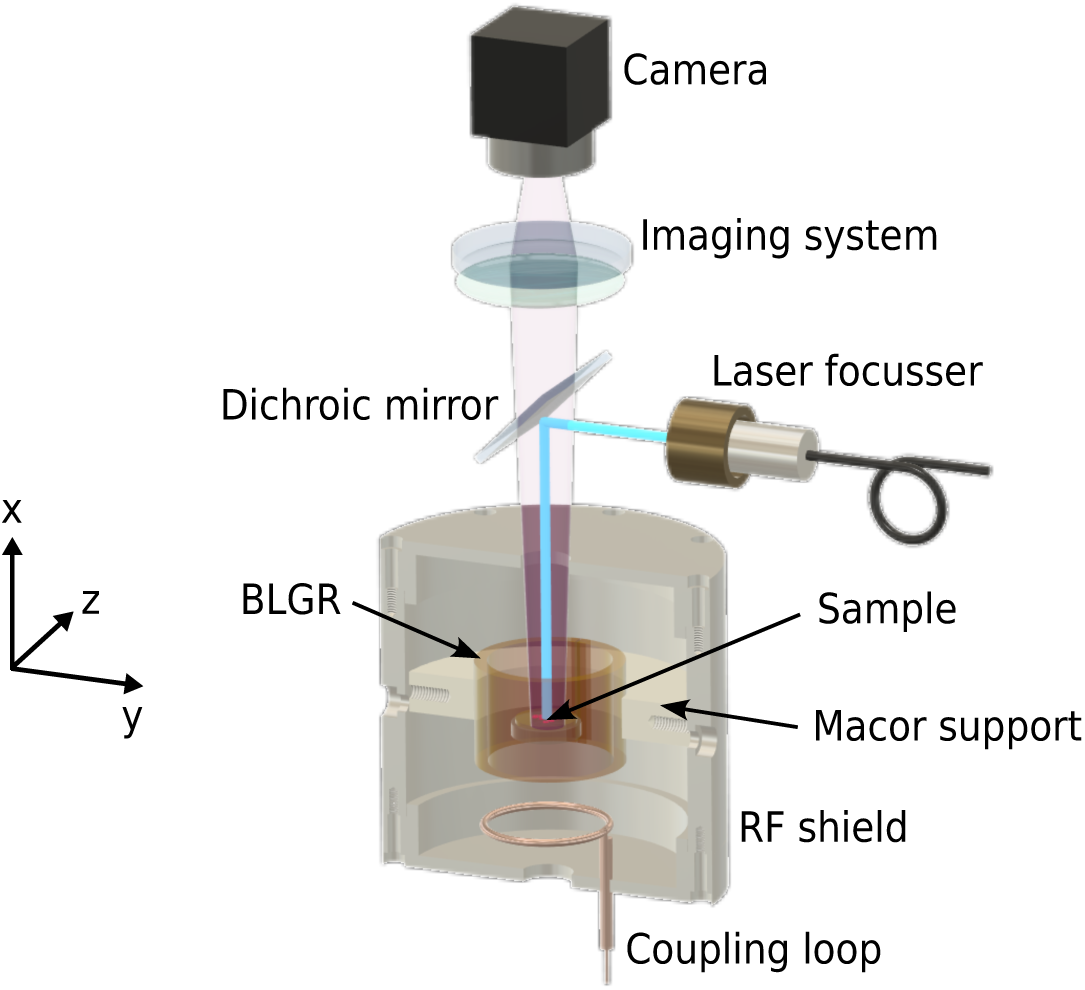
Experimental setup (not to scale). The sample is positioned near the center of a bridged loop-gap resonator (BLGR). Two pairs of Helmholtz coils (not shown) generate static magnetic fields either parallel (*B*_0 ∥_ along *x*) or perpendicular (*B*_0 ⊥_ along *z*) to the RF field direction. A dichroic mirror allows delivery of excitation light at 440, 520, or 561 nm to the sample and transmission of fluorescence to the imaging system and camera.

**Figure S2:**
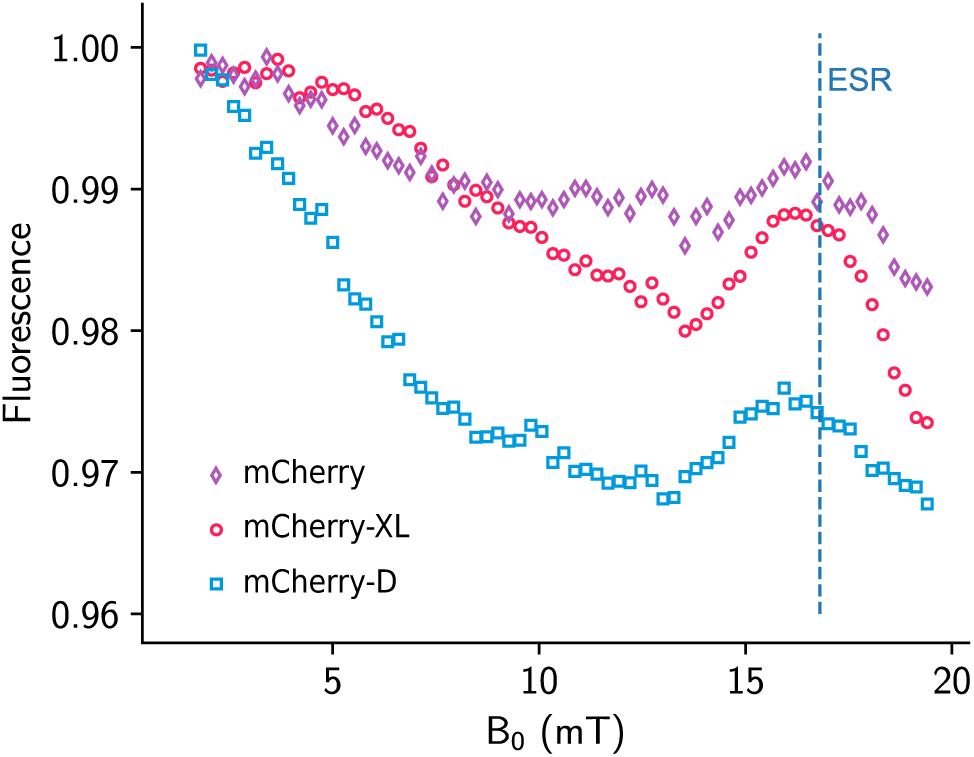
MFEs and RYDMR with mCherry variants. Data points are fluorescence measurements at various values of the static magnetic field *B*_0_ for mCherry (purple diamonds), mCherry-XL (red circles), and mCherry-D (blue squares), in the presence of FMN. The RF field is at 470 MHz, with amplitude *B*_1_ = 0.1 mT. The dashed blue line indicates the magnetic field required for ESR at 470 MHz for a *g* = 2 electron spin.

**Figure S3:**
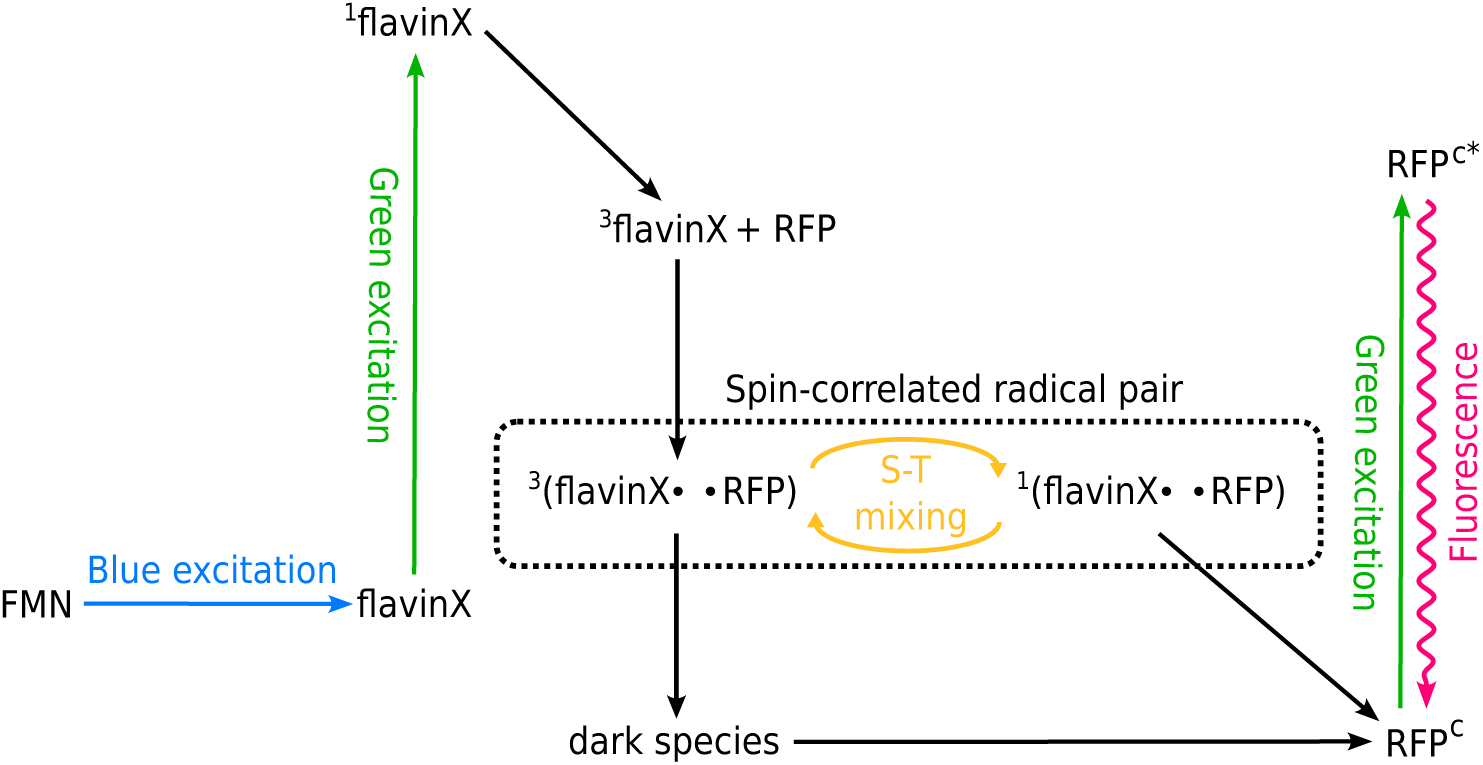
Proposed radical pair reaction scheme for the RFP-flavin system. A two-step process is involved: FMN photoexcitation driven by blue light and a radical pair reaction involving the FMN photoproduct (flavinX) and the RFP. RFP^*c*^ is a representative of the RFP chromophore. After photoexcitation with green light, a spin-correlated RP is formed. Hyperfine interactions drive coherent singlet (*S*) to triplet (*T*) mixing which is sensitive to magnetic fields. The *S* state can undergo a rapid reverse reaction returning the RFP to its molecular ground state. From *T*, the RP can be converted to the *S* state or decay, resulting in formation of a nonfluorescent version of the RFP (dark species), that can decay back to the RFP ground state on much slower time scales. Application of an external magnetic field results in reduced decay from the *S* state and a corresponding reduction in the population of ground state RFP molecules. This results in a reduction in fluorescence.

**Figure S4:**
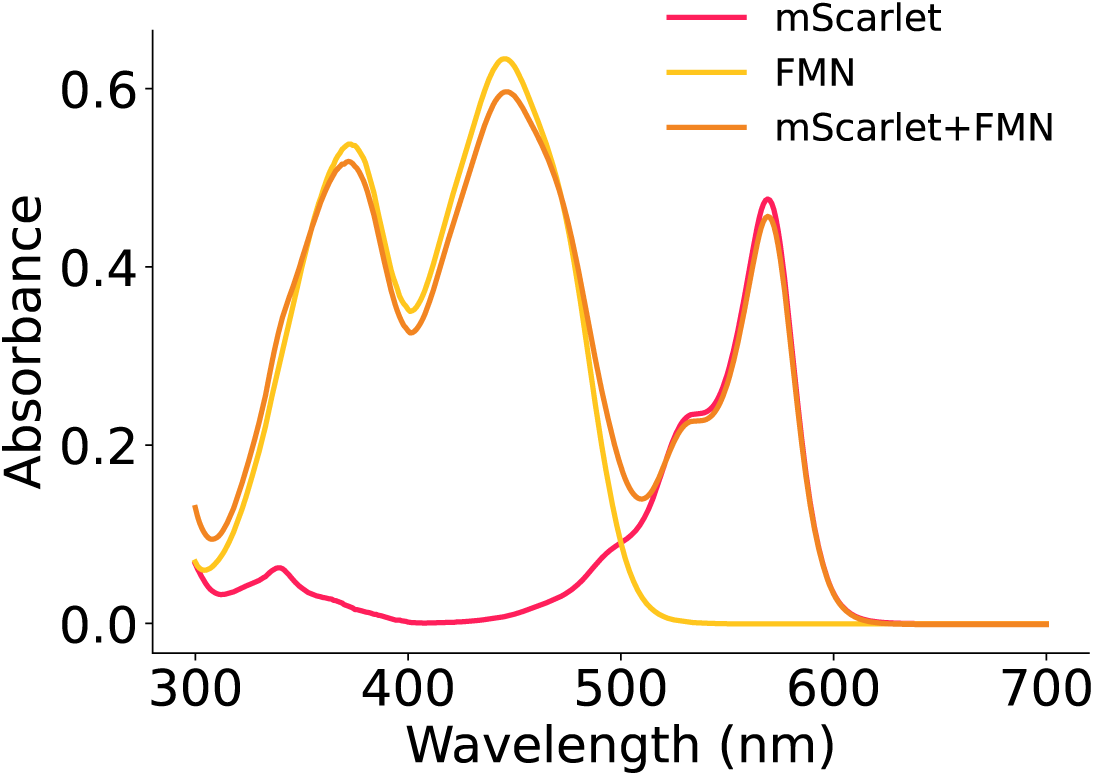
Steady-state absorption spectra for the mixture of mScarlet and FMN, as well as for FMN and mScarlet individually. Concentrations of FMN and mScarlet were about 500 µM and 50 µM, respectively.

**Figure S5:**
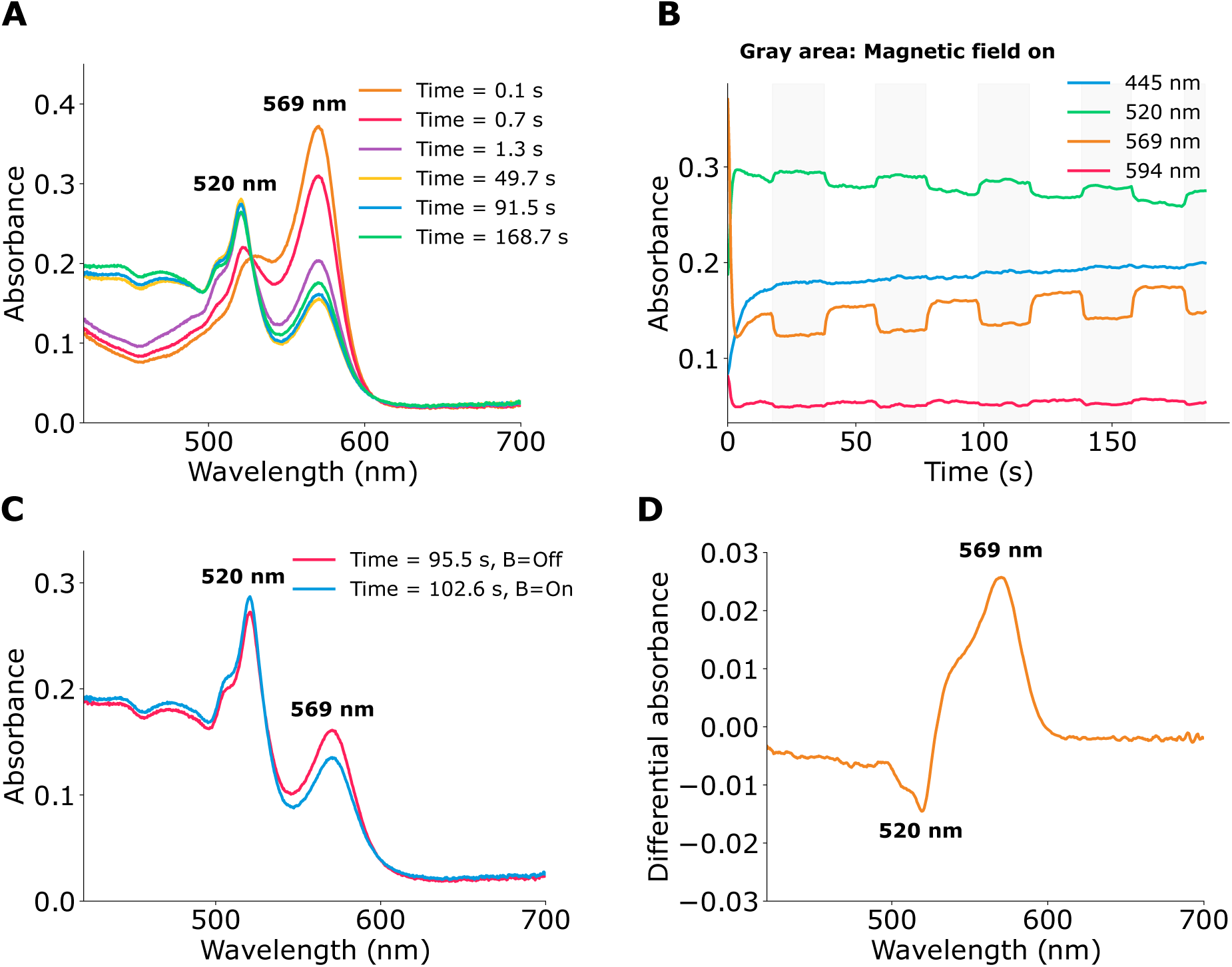
Absorption spectra and magnetically modulated absorption from a solution of mScarlet and FMN. The sample is initially prepared by photoexcitation at 440 nm. (**A**) Absorption from 420 to 700 nm at various durations of 520 nm light exposure. No magnetic field is present in the selected time traces. (**B**) Absorption at various wavelengths as a function of the duration of 520 nm light exposure.White (gray) regions indicate periods when the magnetic field is off (on). (**C**) Absorption with and without the magnetic field. (**D**) Difference in absorption between the traces shown in **C**.

**Figure S6:**
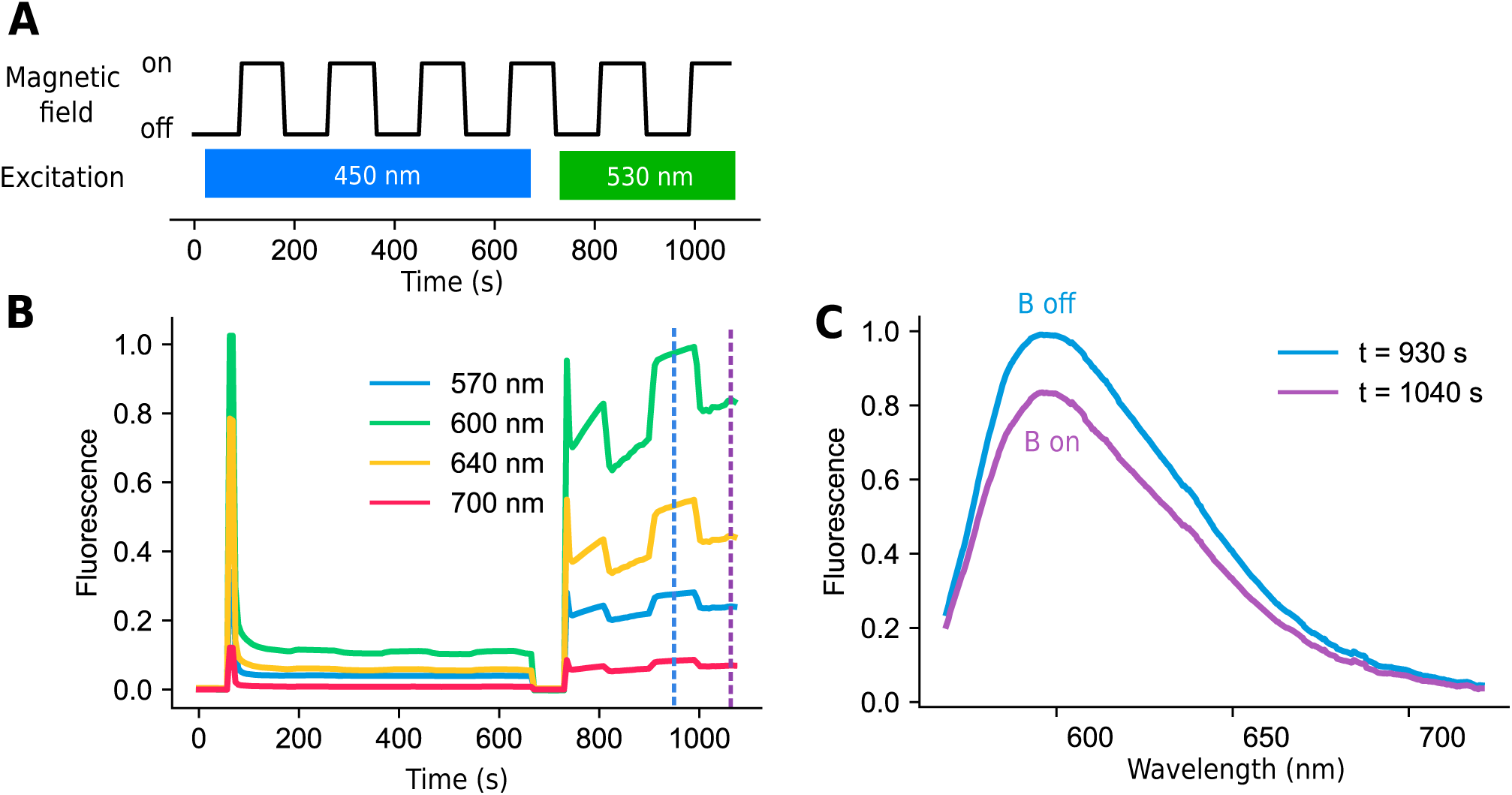
Spectrally-resolved fluorescence magnetic field effects from a solution of 50 µM purified mScarlet-I and 300 µM FMN. (**A**) Timing diagram. The magnetic field is switched between 0 and ∼ 100 mT. (**B**) Spectrally-resolved fluorescence as a function of time following the sequence shown in **A**. Emission spectra over the range 570 to 720 nm with the magnetic field off (t = 930 s) and on (t = 1040 s), indicated by dashed vertical lines in **B**.

**Figure S7:**
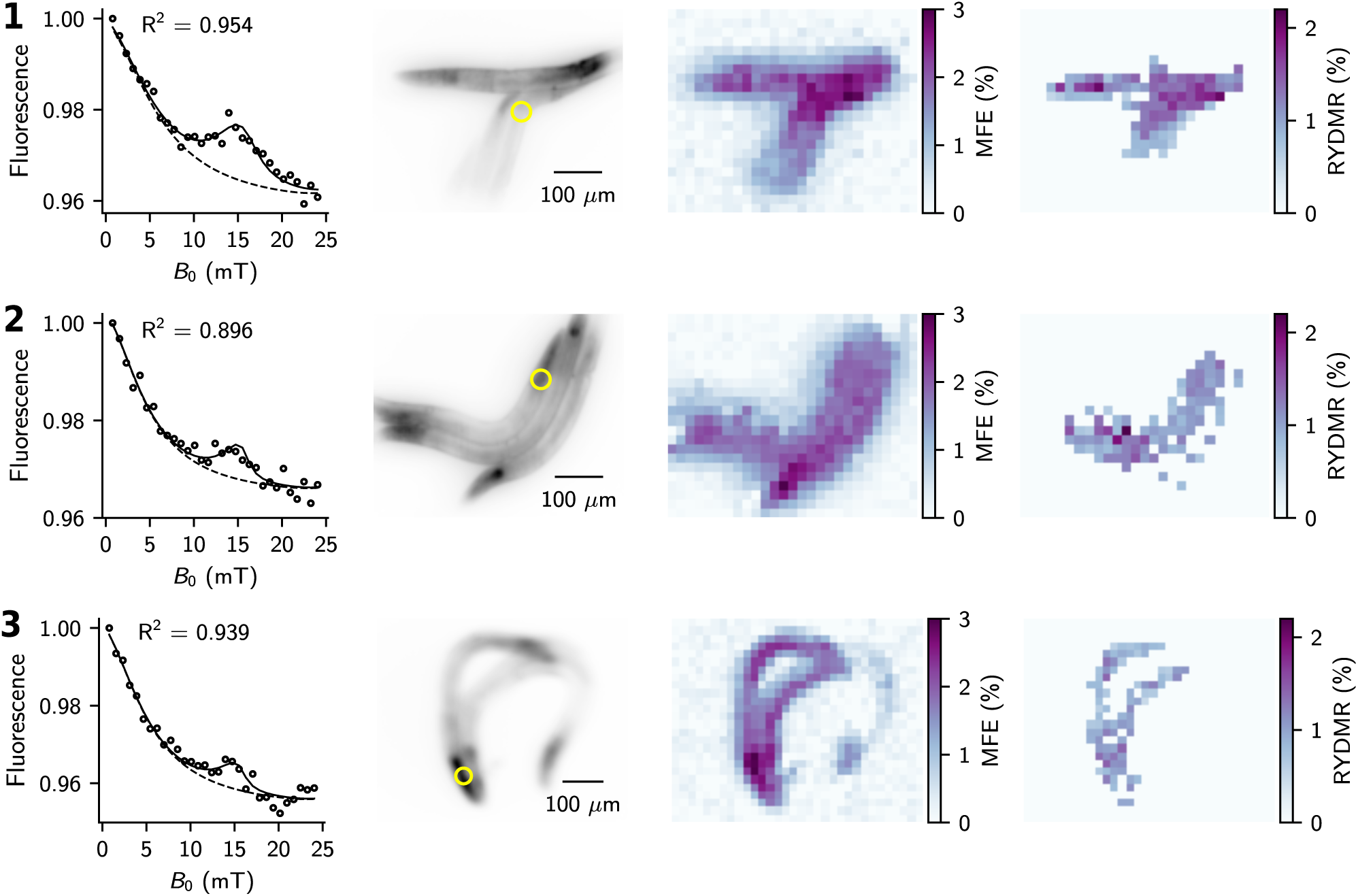
RYDMR from *C. elegans* strain WBM1143. Each row corresponds to a different sample with *B*_1_ = 0.1 mT. Column 1, Integrated fluorescence from within the yellow circles shown in the mScarlet fluorescence images in Column 2 at various values of *B*_0_. Data points are given by black circles. Solid black lines are fits of Eq. S1 to the data. Dashed black lines are estimates of the fluorescence with no RF. Column 3, Spatial distribution of the MFE over the image shown in Column 2 obtained by fitting Eq. S1 to the mean fluorescence from 19 × 19 *μ*m blocks. Column 4, Spatial distribution of RYDMR amplitudes. Only values where *R*^2^ > 0.6 and *p*_F_ < 0.01 from an F test (see SI text) are shown.

**Figure S8:**
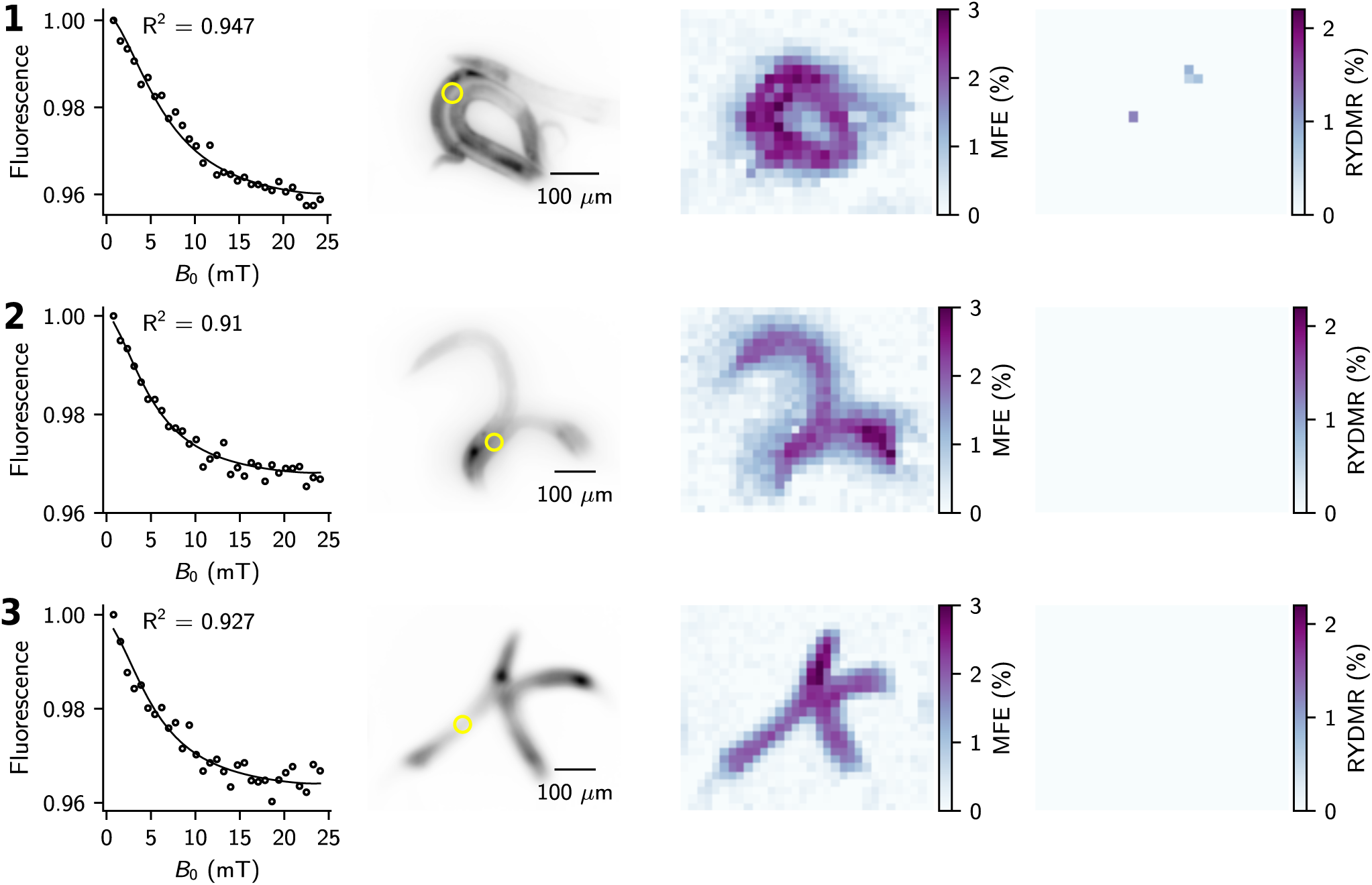
RYDMR from *C. elegans* expressing mScarlet in all cells. Same as for S7, but with *B*_1_ = 0

**Figure S9:**
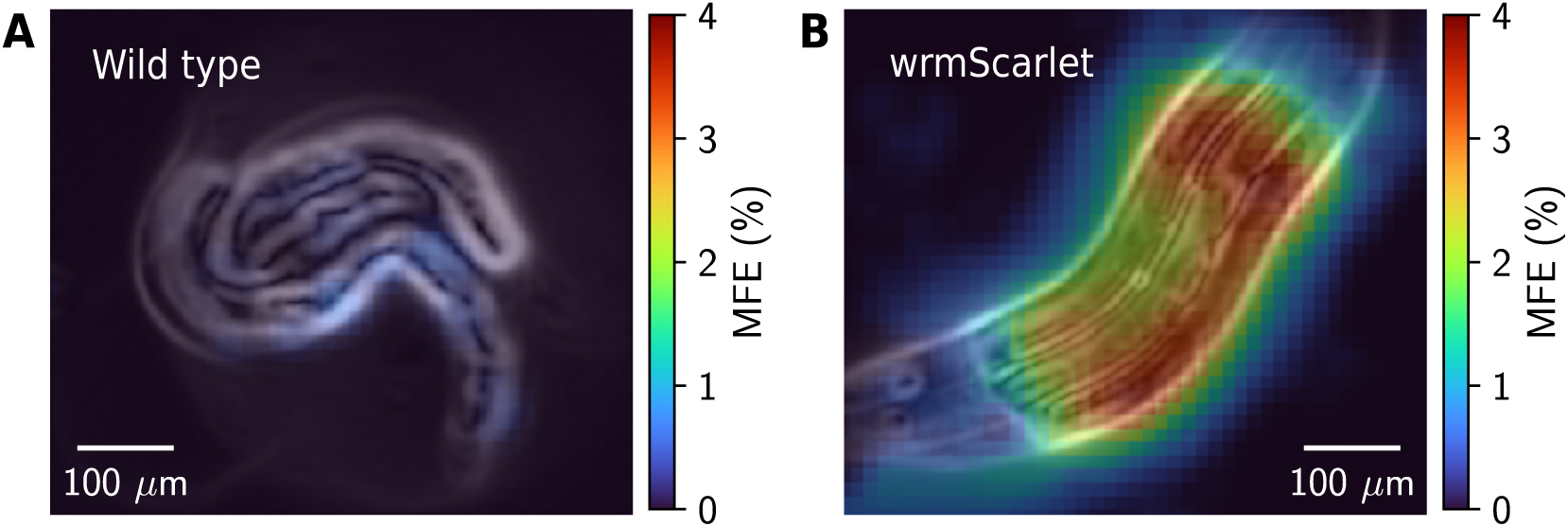
MFEs in fluorescence measured from (**A**) wild-type *C. elegans* and (**B**) *C. elegans* expressing mScarlet in all cells. Each panel shows a color map of the MFE overlaid on an edge map derived from fluorescence images of the nematodes. For these images, fluorescence was collected using a 550 nm long-pass emission filter.

**Figure S10:**
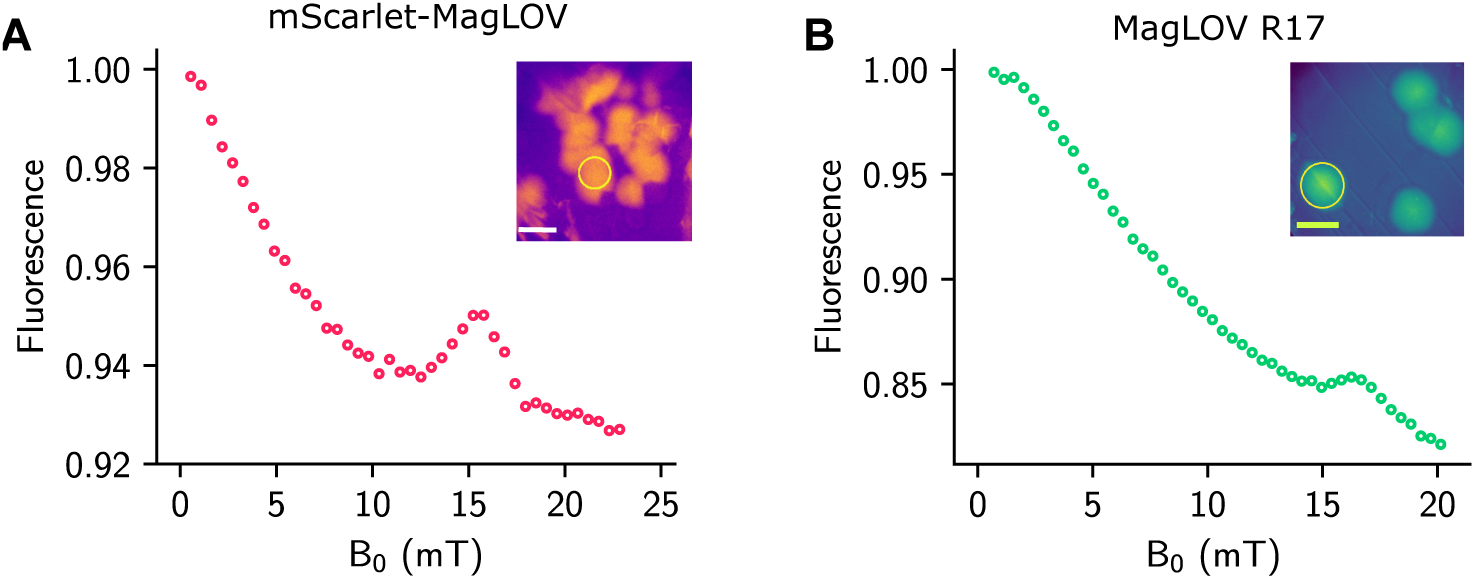
Fluorescence-detected magnetic resonance from *E. coli* colonies expressing mScarlet-MagLOV fusion (**A**) and MagLOV (**B**). Data points are integrated fluorescence from within the regions indicated the yellow circles shown in the insets as *B*_0_ is varied.

## Notes

### Competing Interest Statement

Andrew York and Maria Ingaramo are listed as inventors in U.S. Provisional Patent Application No. 63/568,263, entitled “MUTANT ASLOV2 DOMAINS AND USES THEREOF.”

## References and Notes

1. S. J. Kubarev, E. A. Pshenichnov, The effect of high frequency magnetic fields on the recombination of radicals. Chemical Physics Letters 28 (1), 66–67 (1974).

2. O. A. Anisimov, V. M. Grigoryants, V. K. Molchanov, Y. N. Molin, Optical detection of ESR absorption of short-lived ion-radical pairs produced in solution by ionizing radiation. Chemical Physics Letters 66 (2), 265–268 (1979).

3. U. E. Steiner, T. Ulrich, Magnetic field effects in chemical kinetics and related phenomena. Chemical Reviews 89 (1), 51–147 (1989).

4. B. Leberecht, et al., Upper bound for broadband radiofrequency field disruption of magnetic compass orientation in night-migratory songbirds. Proceedings of the National Academy of Sciences 120 (28), e2301153120 (2023).

5. H. Zadeh-Haghighi, C. Simon, Magnetic field effects in biology from the perspective of the radical pair mechanism. Journal of the Royal Society Interface 19 (193) (2022).

6. P. J. Hore, Spin chemistry in living systems. National Science Review 11 (9), nwae126 (2024).

7. A. Hoff, Magnetic field effects on photosynthetic reactions. Quarterly Reviews of Biophysics 14 (4), 599–665 (1981).

8. S. G. Boxer, C. E. Chidsey, M. G. Roelofs, Magnetic field effects on reaction yields in the solid state: an example from photosynthetic reaction centers. Annual Review of Physical Chemistry 34 (1), 389–417 (1983).

9. T. Miura, K. Maeda, T. Arai, Effect of Coulomb interaction on the dynamics of the radical pair in the system of flavin mononucleotide and hen egg-white lysozyme (HEWL) studied by a magnetic field effect. The Journal of Physical Chemistry B 107 (26), 6474–6478 (2003).

10. E. W. Evans, et al., Sensitive fluorescence-based detection of magnetic field effects in photoreactions of flavins. Phys. Chem. Chem. Phys. 17, 18456–18463 (2015).

11. K. B. Henbest, et al., Magnetic-field effect on the photoactivation reaction of *Escherichia coli* DNA photolyase. Proceedings of the National Academy of Sciences 105 (38), 14395–14399 (2008).

12. T. J. Zwang, E. C. Tse, D. Zhong, J. K. Barton, A compass at weak magnetic fields using thymine dimer repair. ACS Central Science 4 (3), 405–412 (2018).

13. K. Maeda, et al., Magnetically sensitive light-induced reactions in cryptochrome are consistent with its proposed role as a magnetoreceptor. Proceedings of the National Academy of Sciences 109 (13), 4774–4779 (2012).

14. J. Xu, et al., Magnetic sensitivity of cryptochrome 4 from a migratory songbird. Nature 594 (7864), 535–540 (2021).

15. M. K. Bowman, et al., Magnetic resonance spectroscopy of the primary state, P^F^, of bacterial photosynthesis. Proceedings of the National Academy of Sciences 78 (6), 3305–3307 (1981).

16. J. R. Norris, et al., Magnetic characterization of the primary state of bacterial photosynthesis. Proceedings of the National Academy of Sciences 79 (18), 5532–5536 (1982).

17. K. Moehl, E. Lous, A. Hoff, Low-power, low-field RYDMAR of the primary radical pair in photosynthesis. Chemical Physics Letters 121 (1-2), 22–27 (1985).

18. R. Y. Tsien, The green fluorescent protein. Annual Review of Biochemistry 67 (1), 509–544 (1998).

19. M. Chattoraj, B. A. King, G. U. Bublitz, S. G. Boxer, Ultra-fast excited state dynamics in green fluorescent protein: multiple states and proton transfer. Proceedings of the National Academy of Sciences 93 (16), 8362–8367 (1996).

20. D. S. Bindels, et al., mScarlet: A bright monomeric red fluorescent protein for cellular imaging. Nature Methods 14 (1), 53–56 (2017).

21. N. C. Shaner, et al., Improved monomeric red, orange and yellow fluorescent proteins derived from Discosoma sp. red fluorescent protein. Nature Biotechnology 22 (12), 1567–1572 (2004).

22. S. Mukherjee, et al., Directed evolution of a bright variant of mCherry: suppression of nonradiative decay by fluorescence lifetime selections. The Journal of Physical Chemistry B 126 (25), 4659–4668 (2022).

23. S. El Mouridi, et al., Reliable CRISPR/Cas9 genome engineering in Caenorhabditis elegans Using a single efficient sgRNA and an easily recognizable phenotype. G3 Genes|Genomes|Genetics 7 (5), 1429–1437 (2017).

24. C. G. Silva-Garcia, et al., Single-copy knock-in loci for defined gene expression in Caenorhabditis elegans. G3 Genes|Genomes|Genetics 9 (7), 2195–2198 (2019).

25. M. Ono, et al., L-band ESR spectrometer using a loop-gap resonator for in vivo analysis. Chemistry Letters 15 (4), 491–494 (1986).

26. W. N. Hardy, L. A. Whitehead, Split-ring resonator for use in magnetic resonance from 200–2000 MHz. Review of Scientific Instruments 52 (2), 213–216 (1981).

27. R. F. Hayward, et al., Magnetic control of the brightness of fluorescent proteins (2024), doi:10.5281/zenodo.8137174.

28. A. V. Koptyug, V. O. Saik, O. A. Animisov, Y. N. Molin, Spin-locking in concentration-narrowed OD ESR spectra. Chemical Physics 138 (1), 173–178 (1989).

29. E. F. McCord, R. R. Bucks, S. G. Boxer, Laser chemically induced dynamic nuclear polarization study of the reaction between photoexcited flavins and tryptophan derivatives at 360 MHz. Biochemistry 20 (10), 2880–2888 (1981).

30. S. Stob, R. Kaptein, Photo-CIDNP of the amino acids. Photochemistry and photobiology 49 (5), 565–577 (1989).

31. T. Mani, Molecular qubits based on photogenerated spin-correlated radical pairs for quantum sensing. Chemical Physics Reviews 3 (2) (2022).

32. H. Lee, N. Yang, A. E. Cohen, Mapping nanomagnetic fields using a radical pair reaction. Nano Letters 11 (12), 5367–5372 (2011).

33. E. Gintzon, in Microwave Measurements (McGraw-Hill Book Co., New York), p. 448 (1957).

34. J. H. Freed, D. S. Leniart, J. S. Hyde, Theory of saturation and double resonance effects in ESR Spectra. III. RF coherence and line shapes. The Journal of Chemical Physics 47 (8), 2762–2773 (1967).

35. J. R. Taylor, An Introduction to Error Analysis: The Study of Uncertainties in Physical Measurements (University Science Books) (1997).

36. P. Manna, Development and Characterization of Improved Red Fluorescent Protein Variants, Ph.D. thesis, University of Colorado at Boulder, Boulder, CO (2017).

37. S. Mukherjee, Spectroscopic Evaluation of Excited State Depopulation in Red Fluorescent Proteins Developed Using Fluorescence Lifetime Selections, Ph.D. thesis, University of Colorado at Boulder, Boulder, CO (2022).

38. S. Brenner, The genetics of Caenorhabditis elegans. Genetics 77 (1), 71–94 (1974).

39. P. D. Berger, R. E. Maurer, G. B. Celli, Introduction to Experimental Design. Experimental Design: With Application in Management, Engineering, and the Sciences. pp. 1–19 (2018).

40. N. Ikeya, J. R. Woodward, Cellular autofluorescence is magnetic field sensitive. Proceedings of the National Academy of Sciences 118 (3), e2018043118 (2021).

